# The histone demethylase KDM6B fine-tunes the host response to *Streptococcus pneumoniae*

**DOI:** 10.1101/757906

**Authors:** Michael G. Connor, Tiphaine Marie-Noelle Camarasa, Emma Patey, Orhan Rasid, Laura Barrio, Caroline M. Weight, Daniel P. Miller, Robert S. Heyderman, Richard J. Lamont, Jost Enninga, Melanie A. Hamon

## Abstract

*Streptococcus pneumoniae* is a natural colonizer of the human upper respiratory tract and an opportunistic pathogen. After colonization, bacteria either remain in the human upper respiratory tract, or may progress to cause pneumococcal disease. Although epithelial cells are among the first to encounter pneumococci, the cellular processes and contribution of epithelial cells to the host response are poorly understood. Here, we show a *S. pneumoniae* serotype 6B ST90 strain, which does not cause disease in a murine infection model, induces a unique NF-κB signature response distinct from an invasive disease causing isolate of serotype 4 (TIGR4). This signature is characterized by activation of p65 (RelA) and requires a histone demethylase, KDM6B. At the molecular level, we show that interaction of the 6B strain with epithelial cells leads to chromatin remodeling within the IL-11 promoter in a KDM6B dependent manner, where KDM6B specifically demethylates histone H3 lysine 27 di-methyl. Chromatin remodeling of the IL-11 locus facilitates p65 access to three NF-κB sites, which are otherwise inaccessible when stimulated by IL-1β or TIGR4. Finally, we demonstrate through chemical inhibition of KDM6B, with GSK-J4 inhibitor, and through exogenous addition of IL-11 that the host responses to 6B ST90 and TIGR4 strains can be interchanged both *in vitro* and in a murine model of infection *in vivo*. Our studies hereby reveal how a chromatin modifier governs cellular responses during infection.

## Introduction

*Streptococcus pneumoniae* (the pneumococcus), a heterogeneous species comprised of over 90 serotypes, naturally colonizes the upper respiratory tract of humans, and is also an opportunistic pathogen ^1-6^. Globally, disease attributed to *S. pneumoniae* infection is a priority due to potential lethality resulting from pneumonia, sepsis, or meningitis ^4,7^. Importantly, colonization of the human upper respiratory tract is the natural reservoir for this obligate pathobiont. Here, pneumococcus either remains in the nasopharynx and is eventually cleared, or may progress to cause a disease state ^2,3,5,8,9^.

At these initial stages of pneumococcal-epithelial interaction, the pneumococcus interacts with the host nasopharyngeal epithelial barrier and the innate immune system. Recent insights using the experimental human pneumococcal carriage (EHPC) model have highlighted the essential role of NF-κB driven responses for susceptibility, pathogenesis and transmission of pneumococcus ^8,10-13^. However, it is unclear how these cellular processes are shaped at the molecular level, and how they affect the potential progression to pneumococcal disease ^14^.

NF-κB is a master transcriptional regulator of both pro- and anti-inflammatory host responses ^15-25^. Briefly, NF-κB is comprised of multiple subunits that form hetero- or homodimers, of which the best characterized subunit is p65 (RelA) ^26,27^. Activation of p65 occurs through posttranslational modifications (PTMs), such as phosphorylation of serine 536, in response to cellular sensing of inflammatory stimuli (i.e. lipopolysaccharide (LPS) or interleukin 1 beta (IL-1β)) ^27^. Activated p65 binds to a kappa-binding consensus sequence sites within the nucleus to initiate transcription of NF-κB dependent genes ^28^. However, cellular signaling alone is not enough, as a full NF-κB response also requires chromatin remodeling at the targeted inflammatory gene loci ^15-25^.

Chromatin is a highly ordered structure of DNA wrapped around histone proteins. Chromatin dynamically shifts between open (euchromatin), and closed (heterochromatin) states, which influence gene accessibility and transcription ^29-33^. Switching between these two states is the result of chromatin remodeling enzymes/complexes reading, writing and erasing PTMs on histone tails. The enzymes regulating histone PTMs have been identified, and shown to play important roles in transcriptional responses during cellular signaling events, such as NF-κB responses ^15,34^. One of these enzymes is KDM6B (JMJD3), a histone demethylase, associated with NF-κB. KDM6B belongs to the Jumonji C-domain family (JMJD) of histone demethylases, of which KDM6B is the only member expressed universally outside of embryonic development ^23,35^. Primarily through peptide studies, KDM6B is thought to target the repressive histone marks, lysine 27 tri-methyl (H3K27me3) and di-methyl (H3K27me2) ^23,36-38^. To date, mounting evidence, mainly in macrophages, suggests KDM6B is essential for modulating inflammatory gene expression during wound healing, upon LPS stimulation and for immunological tolerance to anthrax toxin ^19-22,39^. However, the role of KDM6B in bacteria induced host responses has not been studied.

Herein, we demonstrate a pneumococcal isolate of serotype 6B ST90 CC156 lineage F, which in a murine infection model is contained to the upper respiratory tract and does not cause invasive disease, specifically activates a unique NF-κB signature distinct from the invasive disease causing serotype 4 isolate TIGR4. This signature includes upregulation of KDM6B and the cytokine IL-11 in a human epithelial cell culture model. We show that the promoter of IL-11 is remodeled upon challenge with the 6B strain via KDM6B demethylation of H3K27me2, which allows p65 binding at three NF-κB sites upstream of the IL-11 transcription start site. We establish the importance of this process in regulating epithelial cell integrity and show that, *in vivo* and *in vitro*, chemical inhibition of KDM6B causes the 6B strain, which is otherwise non-invasive, to cause disease. Conversely, exogenous addition of recombinant IL-11 during challenge with an invasive TIGR4 isolate leads to better containment of this strain to the upper respiratory tract in a murine model of infection.

## Results

### Serotype 6B ST90 and TIGR4 display different disease outcomes in a murine model of intranasal infection

Using a murine model of infection, we preformed comparative intranasal challenge studies of pneumococcal isolates TIGR4 (serotype 4) and 6B ST90 CC156 lineage F (serotype 6B; hereafter referred to as 6B). Infection of male mice with 3 - 4×10^6^ CFU (Sup. Fig. 1A) of TIGR4 induced rapid weight loss and morbidity, within 4 days post-challenge (Fig. 1A & B). In contrast, the 6B isolate did not cause symptomatic pneumococcal disease, as measured by weight loss and survival, even at an infection dose one log higher than that of TIGR4 (Fig 1A & B). Since the murine host outcomes were drastically different, we chose to study the molecular processes in more detail using the 6B and TIGR4 strains.

**Figure 1:**
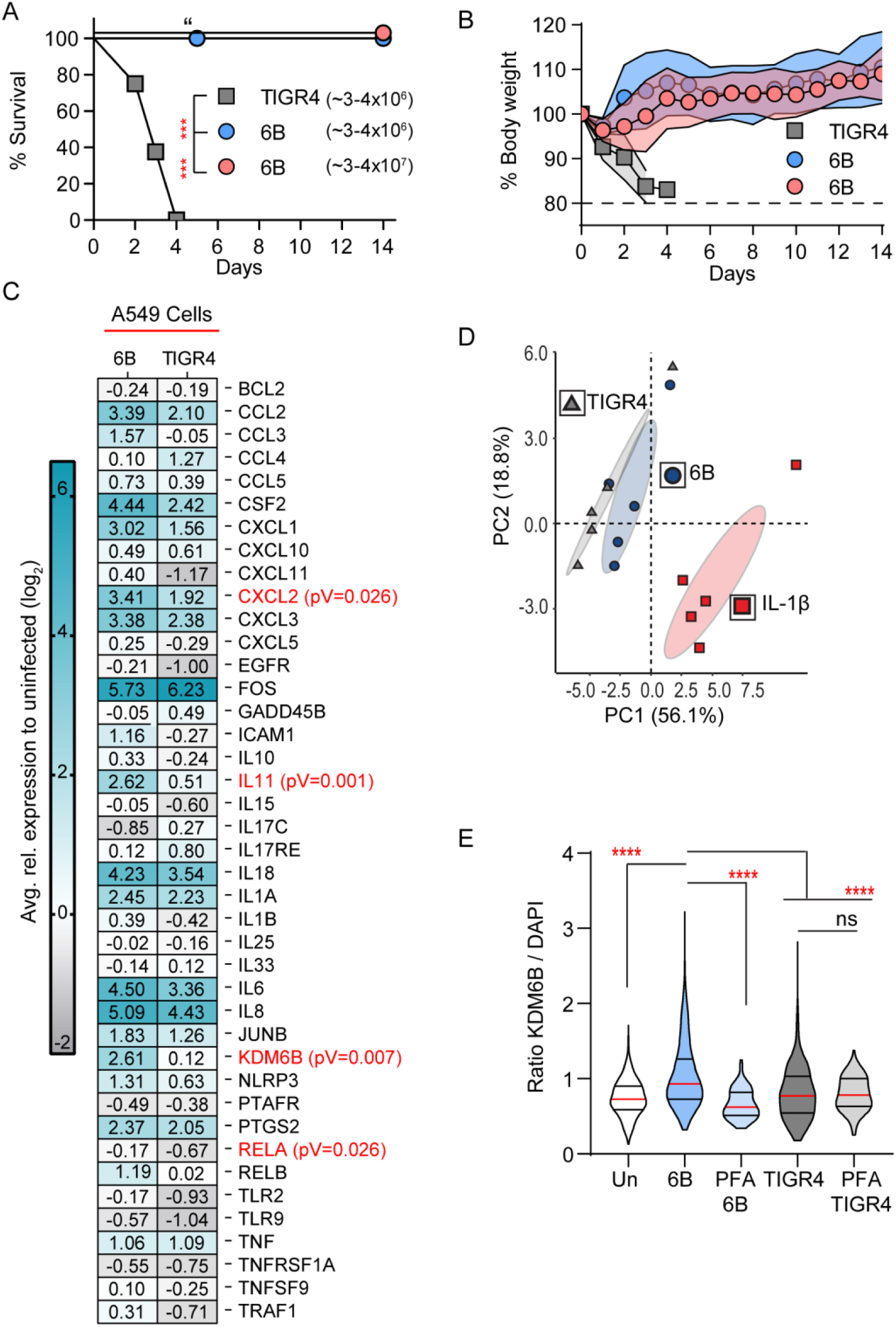
Inflammatory signature of serotype 6B ST90 CC156 lineage F strain. C57B6 mice (8-9weeks) were challenged intranasally with TIGR4 (3 – 4×10^6^ CFU; n=8), 6B (3 - 4×10^6^ CFU; n=6), or 6B (3 - 4×10^7^ CFU; n=6). Survival and weight monitored for 14 days post-infection. A) Percent survival curve. Mantel-Cox test for significance, ***= pV≤0.001. “= animal euthanized but did not succumb to disease. B) Percent initial body weight. Solid lines per group are Std. Dashed line indicates 20% weight loss threshold. C) RT-PCR validation of inflammatory genes from cells collected 2 hrs post-challenge with either TIGR4 (MOI 20) or 6B (MO20). Heat map represents fold change to uninfected condition (n=5). Multiple T-Test 6B to TIGR4 with significant genes in red. D) Principal component analysis of inflammatory RT-PCR panel (n=5) comparing IL-1β (red), TIGR4 (gray) and 6B (blue). Bioplot of the mean centered and log_2_ transformed expression data using the first two components and 95% confidence ellipses around each group. E) Quantification of immunofluorescence microscopy of A549 cells 2 hrs post-challenge with either TIGR4 (MOI 20), 6B (MOI 20) or paraformaldehyde fixed 6B or TIGR4 (MOI 20) for nuclear staining of KDM6B normalized to DAPI (n=4; 30-100 cells per replicate and group). Violin plots with median denoted in red. One way-ANOVA non-parametric Kruskal-Wallis test with Dunn’s multiple comparison post-hoc test, ****= pV≤0.0001.

### Serotype 6B ST90 actively induces a unique cellular response

To begin dissecting the host processes differentially regulated by the strains 6B and TIGR4, we completed an exploratory microarray of human A549 epithelial cells 2 hrs post-challenge (Sup. Fig 1B). In comparison with uninfected cells, 6B differentially influenced 388 transcripts (200 upregulated and 188 downregulated); whilst TIGR4 modulated the expression of 1,205, (143 upregulated and 1,062 downregulated) (Sup. Table 1). Strikingly by 2 hrs post challenge a larger proportion of the total genes within the 6B dataset contained NF-κB binding sites in contrast to TIGR4 (12% by 6B vs. 3% by TIGR4; Sup. Fig. 1C). This suggested that early NF-κB regulation of gene transcription influenced the host response to 6B. To test this, we selected a panel of 41 genes, which included genes from the microarray (*IL-11, KDM6B (JMJD3), PTGS2, CXCL8 (IL-8), FOS* and *JUNB*) and 32 direct targets of NF-κB for testing by RT-PCR. Upon A549 epithelial cell challenge with the 6B strain a significant increase in the expression of *CXCL2, IL-11, KDM6B*, and *RELA* in comparison to TIGR4 was observed by 2 hrs (Fig. 1C; Sup. Table 2). This cellular response to the 6B strain was not restricted to A549 type II epithelial cells, but was also observed with nasopharyngeal Detroit 562 and the non-cancer Beas2b epithelial (type II) cell lines as KDM6B and IL-11 clustered across cell lines with elevated expression levels for 6B in comparison to TIGR4 (Sup Fig. 1D; Sup. Table 2). We further cross-compared the same panel of genes upon IL-1β stimulation (Sup. Table 2), a known pro-inflammatory stimulus which activates p65 (RelA). Using the relative expression data obtained from RT-PCR of A549 cells, we performed a principal component analysis (PCA) on the expression values for 6B, TIGR4 and IL-1β (Fig. 1D). Comparative analysis of the biplot of the first two components showed three groups (95% confidence ellipses), which accounted for 74.9% of the total variance. Altogether, these data demonstrated 6B was inducing an epithelial cellular response characterized by expression of a subset of inflammatory genes, distinct from both TIGR4 and IL-1β.

To determine if RT-PCR expression results for *KDM6B*, one of the genes differentially expressed at 2 hrs post-challenge, were reflected at the protein level, we performed immunofluorescence staining for KDM6B (Fig. 1E; Rep. images Sup. Fig. 1E). A549 cells were challenged with either live or paraformaldehyde-killed 6B and TIGR4 strains. Two hours post-challenge the nuclear signal intensity of KDM6B to DAPI was quantified. For 6B there was a significant increase in KDM6B (pV≤0.0001) compared to TIGR4 and uninfected cells. We did not see a significant increase in KDM6B signal intensity following challenge with either paraformaldehyde-killed 6B or TIGR4. This showed that increased KDM6B expression is an active process due to pneumococcal-epithelial interaction, as paraformaldehyde fixation not only inactivates pneumococcus, but is known to maintain bacterial morphology including pili, and extracellular polymeric substances, such as capsule ^40^. Additionally, we sampled supernatants from Detroit 562 cells 6 hrs post challenge with either 6B or TIGR4, and observed a modest increase in detectable IL-11 (Sup. Fig. 1F).

Since KDM6B is part of a larger Jumonji C-domain (JMJD) histone demethylase family, we determined if 6B challenge upregulated additional JMJD demethylases, or was specific to KDM6B. We tested three additional JMJD demethylases (KDM7A, KDM8, and KDM6A (UTX)), and an unrelated methyltransferase (EHMT2) by RT-PCR. KDM6B was the only one to be significantly upregulated by 6B (Sup. Fig. 1G). Together these results show 6B induces a differential transcriptional response in epithelial cells characterized, in part, by upregulation of KDM6B and the cytokine IL-11.

### 6B cellular response requires p65 activation and catalytically active KDM6B

A hallmark of NF-κB dependent gene induction is the activation of p65 via phosphorylation at serine 536 (S536) ^26^. Thus, we tested if phosphorylation of p65 occurred during 6B challenge. Whole cell lysates were collected 2 hrs post-challenge from HeLa GFP-p65 stable cell line, and were immunoblotted for p65 phosphorylation at S536 (Fig. 2A). In comparison to uninfected cells, both IL-1β (positive control) and 6B induced p65 phosphorylation of S536 (pV≤0.001) whereas TIGR4 did not (Fig. 2B). To determine whether p65 activation by 6B was required for *IL-11* and *KDM6B* expression, we used a chemical inhibitor of p65 activation, BAY 11-7082 ^41-43^. A549 cells were pretreated for 3 hrs with 10 µM BAY 11-7082 prior to challenge with 6B, TIGR4 or IL-1β, and gene expression was determined by RT-PCR. BAY 11-7082 treatment did not affect viability or gene expression alone in comparison to untreated cells (Sup. Fig. 2A & B; gray bars). As a control for inhibitor effect, we used the gene *PTGS2*, which is known to require p65 activation for expression. During 6B challenge no significant effect between untreated (no inhibitor) and DMSO (vehicle control) was observed upon the expression of *PTGS2, IL-11* and *KDM6B*, and under the same conditions *IL-11* and *KDM6B* expression remained 2-fold higher on average in comparison to TIGR4 and IL-1β (Fig. 2C; white & light gray bars). Inhibiting p65 activation during 6B, TIGR4 or IL-1β challenge resulted in reduction in *PTGS2* (Fig. 2C; gray bars). The expression of *KDM6B* and *IL-11* were significantly repressed during 6B challenge in the presence of inhibitor (pV≤0.001 and 0.05 respectfully) in comparison to DMSO treated cells (Fig. 2C; gray bars). These data show expression of *IL-11* and *KDM6B* is dependent on p65 activation during 6B challenge of epithelial cells. Previous studies demonstrated KDM6B interacts with p65 for gene activation during keratinocyte wound healing, and ChIP-seq studies found LPS stimulation of macrophages lead to KDM6B regulation of specific inflammatory genes ^20,22^. To determine whether KDM6B had an active role in 6B induced expression of *IL-11* and *KDM6B*, we used GSK-J4, an inhibitor of the catalytic JMJ domain of KDM6B ^44^. As a control, we chose expression of *PTGS2*, as it is associated with KDM6B and not H3K27me3, thus inhibition of the catalytic activity of KDM6B should have no effect upon its expression ^20^. GSK-J4 (10 µM; 24 hrs prior) was used to pretreat A549 cells before challenge with 6B, TIGR4 or IL-1β. GSK-J4 alone had no significant effect on cell viability, or gene expression in comparison to untreated cells, nor did it affect the transcripts of *PTGS2, IL-11* or *KDM6B* in TIGR4 and IL-1β challenged cells (Fig. 2C and Sup. Fig. 2A & B; black bars). Whereas, when the catalytic activity of KDM6B was inhibited, more than a 3-fold loss of expression for both *IL-11* and *KDM6B* was observed during 6B challenge compared to DMSO control (Fig. 2C; black bars). GSK-J4 treatment had no effect upon *PTGS2* expression during 6B challenge, demonstrating KDM6B catalytic activity was specifically required for IL-11 expression (Fig. 2C; black bars).

**Figure 2:**
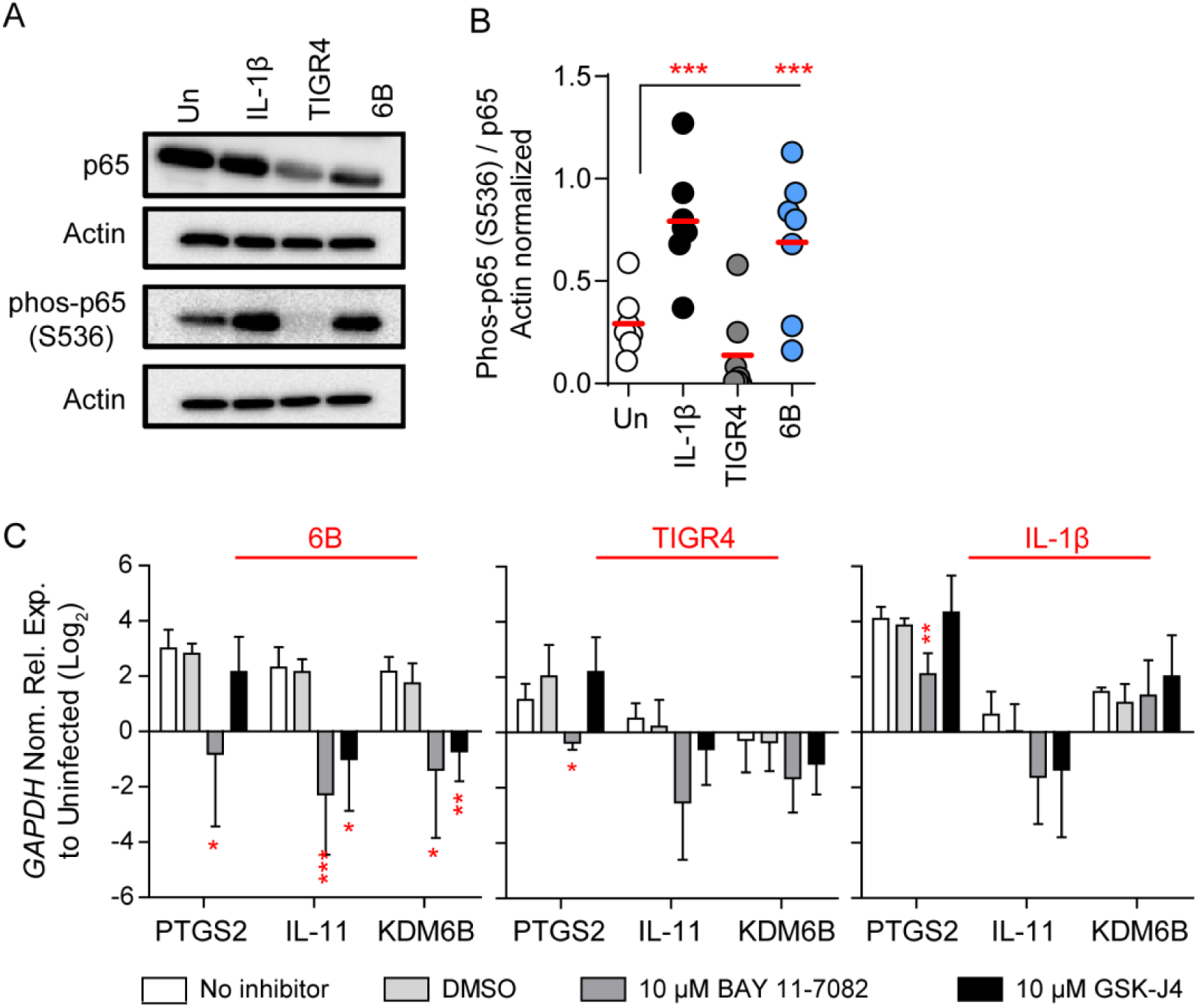
Expression of KDM6B and IL-11 is specific to 6B and requires p65 activation. Immunoblot of stable HeLa p65-GFP expressing cells. Whole cell lysates 2 hrs post-challenge with either IL-1β (10 ng/mL), TIGR4 (MOI 20), or 6B (MOI 20) and probed for p65, phosphorylated p65 at Serine 536 and actin. A) Representative image of immunoblot. B) Actin normalized ratio phos-p65 S536 to total p65 (n=7). Dot plot with mean denoted in red. One way-ANOVA with Tukey’s multiple comparison post-hoc test, **= pV≤0.01, ***= pV≤0.001. C) Total RNA 2 hrs post-challenge with 6B (MOI 20), TIGR4 (MOI 20) or IL-1β was harvested from A549 cells treated with either 10 µM BAY 11-7082, 10 µM GSK-J4, or DMSO vehicle control (n=4 per group). Transcript levels for *PTGS2, IL-11* and *KDM6B* determined by RT-PCR. Bar graph ± Std. Student’s T-Test to untreated, *= pV≤0.05, **= pV≤0.01, ***= pV≤0.001.

### Chromatin is remodeled within the IL-11 promoter upon 6B challenge

Since, KDM6B is p65 dependent, we addressed whether 6B induced p65 recruitment to KDM6B, and if the expression of *IL-11* required KDM6B dependent chromatin remodeling within the IL-11 promoter. We mapped and designed ChIP-qPCR primers to predicted kappa-binding sites within the KDM6B and IL-11 promoter using AliBaba2 software, which curates eukaryotic transcription factor DNA binding motifs from the TRANSFAC^®^ database ^45^. Two kappa-binding sites upstream (−880bp, and −41bp) of the *KDM6B* transcriptional start site (TSS) were found, while three kappa-binding sites upstream (−2,077bp, −774bp, and −406bp) of the *IL-11* TSS, and one site downstream (+83bp) were predicted (Fig. 3A & Sup. Fig. 3C). Herein, we obtained chromatin from A549 cells 2 hrs post-challenge with either 6B, TIGR4, or IL-1β with and without chemical inhibition of the catalytic activity of KDM6B. First, we confirmed p65 recruitment to kappa-bindings sites within the *KDM6B* promoter using ChIP-qPCR (Sup. Fig. 3A & B). We next used ChIP-qPCR to compare the recovery of p65, and KDM6B at these kappa-binding sites within the *IL-11* promoter. During 6B challenge there was a significant (pV≤0.001) recovery of p65 at kappa-binding sites P6 (25%), P3 (20%) and P2 (10%) in contrast to uninfected conditions (Fig. 3B; 6B dark blue; uninfected white). Furthermore, there was 15% recovery of KDM6B across the same kappa-binding sites in cells challenged with 6B (Fig. 3C; 6B dark blue; uninfected white). In contrast, there was no recruitment of p65 or KDM6B to the kappa-binding sites in IL-1β or TIGR4 challenged cells (Sup. Fig. 3D & E). Recruitment of p65 and KDM6B to these kappa-binding sites was abolished in the presence of the GSK-J4 inhibitor (Fig. 3B & C; 6B light blue; uninfected gray). This showed during 6B challenge the promoter of *IL-11* was rearranged in a manner requiring the catalytic activity of KDM6B.

**Figure 3:**
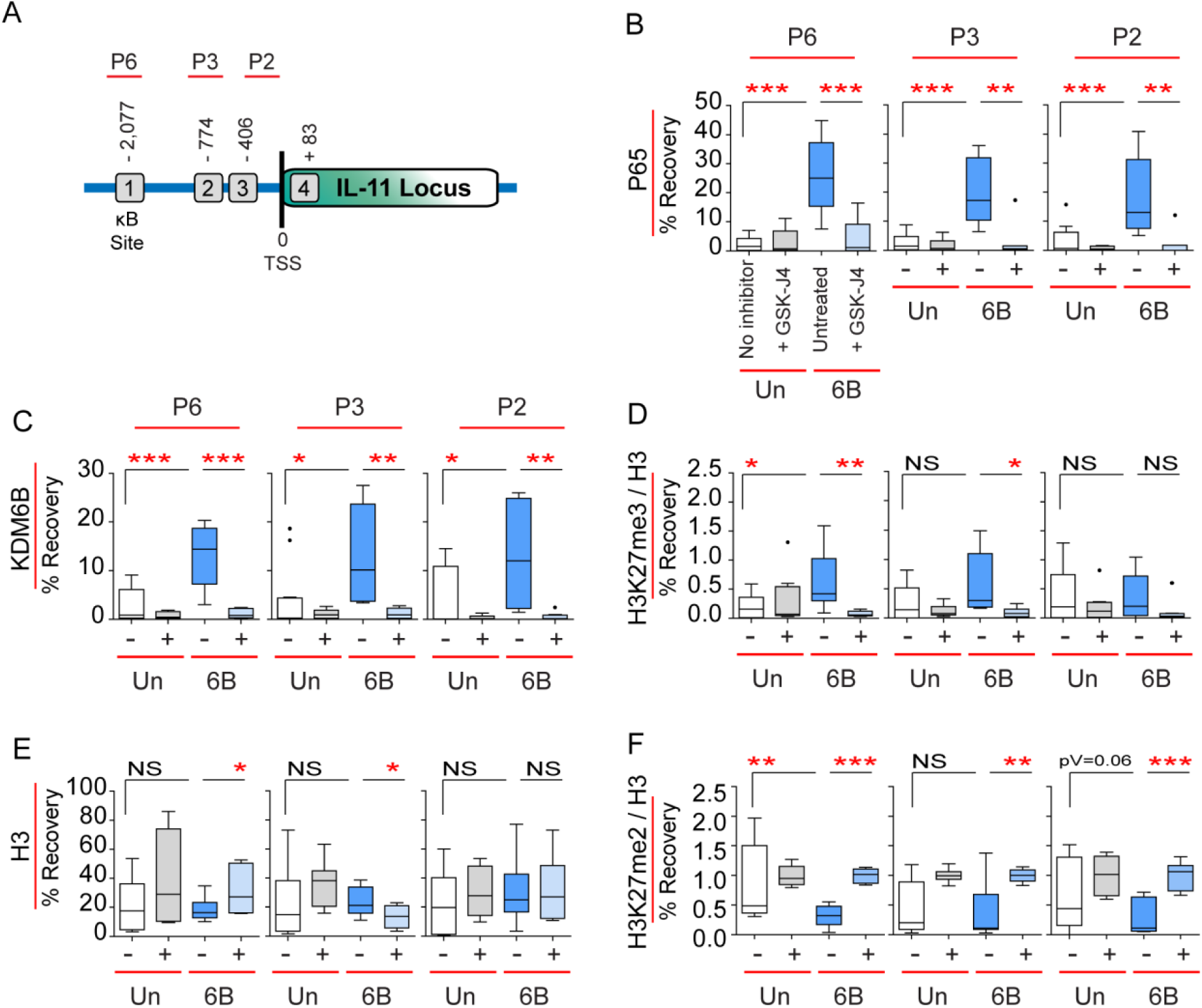
6B induces IL-11 promoter rearrangement. Chromatin was obtained from untreated (blue) and 10 µM GSK-J4 (light blue) treated A549 cells 2 hrs post-challenge with 6B (MOI 20) in comparison to uninfected (untreated white; treated gray). 10 µg chromatin input used for ChIP of p65, KDM6B, H3K27me3 and histone H3 (H3), followed by ChIP-qPCR at locations (P6, P3 & P2) spanning the NF-κB sites upstream of the transcriptional start site (TSS). A) Schematic of IL-11 promoter with ChIP-qPCR primer locations (P6, P3 & P2) and NF-κB sites. B - F) % recovery of input for p65 (B), KDM6B (C), H3K27me3 normalized to H3 (D), H3K27me2 normalized to H3 (F), or H3 (E) bound at P6, P3 & P2 in untreated and GSK-J4 treated samples (n=3 untreated; n=3 GSK-J4 treated). Tukey box and whisker plot with dots representing outliers. Student’s T-Test comparisons for untreated to GSK-J4 treated or 6B infected to uninfected, *= pV≤0.05, **= pV≤0.01, ***= pV≤0.001, ns=not significant.

It has been suggested, mainly through peptide studies, that the enzymatic target of KDM6B is primarily H3K27me3 ^38,46^. Thus, we hypothesized chromatin rearrangement within the *IL-11* promoter was a result of KDM6B demethylation of H3K27me3. We used ChIP-qPCR to determine the levels of H3K27me3 and H3, for nucleosome occupancy, across the three kappa-binding sites within the *IL-11* promoter. Surprisingly, H3K27me3 was not decreased, in fact there was a slight, but significant (pV≤0.05), increase at the P6 kappa-binding site in comparison to unchallenged cells (Fig. 3D; 6B dark blue; uninfected white). There was no enrichment at any kappa-site during IL-1β or TIGR4 challenge (Sup. Fig. 3F). Furthermore, there were no differences in H3 nucleosome distribution at any of the kappa-binding sites between 6B and uninfected cells (Fig. 3E; 6B dark blue; uninfected white), this was also the case for cells challenged with IL-1β or TIGR4 (Sup. Fig. 3G). In the presence of GSK-J4 the increase of H3K27me3 at the P6 and P2 kappa-binding sites was lost in conjunction with a slight but significant increase in H3 nucleosome recovery at P6 (Fig. 3E & D; 6B light blue; uninfected gray). This data showed that during 6B challenge, KDM6B was not demethlyating H3K27me3, and this posttranslational modification seemed to increase across the promoter.

Since our data showed an active role for KDM6B enzymatic activity independent of H3K27me3, we tested another proposed substrate of KDM6B, H3K27me2 ^37^. Our ChIP-qPCR results showed challenge with 6B induced loss of H3K27me2 at the P6 (pV≤0.01) and variable levels at the P3 and P2 sites within the *IL-11* promoter in comparison to uninfected cells (Fig. 3F; 6B dark blue; uninfected white). Strikingly, when KDM6B enzymatic activity was blocked during 6B challenge demethylation of H3K27me2 was significantly inhibited across all kappa-binding sites (Fig. 3F; 6B light blue; uninfected gray).

Together these data show: 1) upon 6B challenge of epithelial cells the promoter of *IL-11* is remodeled through the cooperative role of KDM6B and p65, and 2) KDM6B enzymatic activity is directed toward H3K27me2 and independent of H3K27me3 at these kappa-binding sites.

### KDM6B and IL-11 shapes the cellular response

Since KDM6B plays a role in chromatin remodeling and is necessary for *IL-11* expression, we hypothesized it was controlling expression of the host response to 6B challenge. To address this, we used RT-PCR to test the expression of our panel of 41 genes in the presence of GSK-J4. GSK-J4 significantly dampened expression of *IL-8* (pV≤0.025), *IL-6* (pV≤0.026), *CFS2* (pV≤0.044), *CCL2* (pV≤0.011), *CXCL1* (pV≤0.049), *KDM6B* (pV≤0.033), and *IL-11* (pV≤0.006; Fig. 4A; Sup. Table 3). In parallel, we hypothesized that exogenously adding IL-11, which is regulated by KDM6B, would alter the host response to TIGR4 during infection. Therefore, we tested, by RT-PCR, the expression levels of NF-κB regulated genes during TIGR4 challenge supplemented with exogenous IL-11. Exogenous IL-11 modified the host response to TIGR4, as the expression of 13 transcripts were significantly changed (Fig. 4B; Sup. Table 3). These data suggested that KDM6B and IL-11 are differentially modulating the cellular host response to either 6B or TIGR4 at specific gene loci.

**Figure 4:**
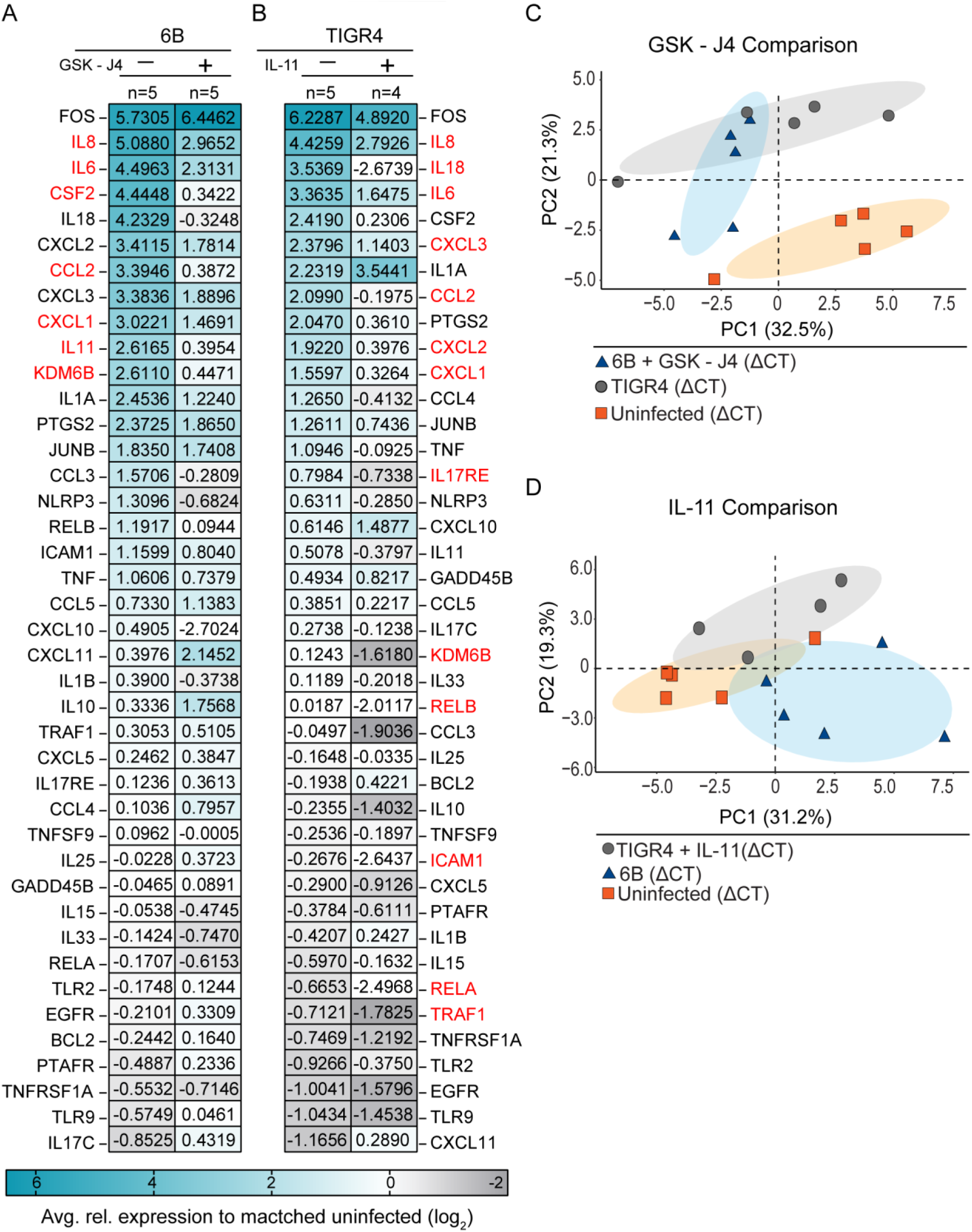
Comparison of GSK-J4 and IL-11 treatment effects upon the host response to 6B and TIGR4. A549 cells untreated or treated with 10 µM GSK-J4 or 100 ng/mL recombinant human IL-11 prior to 2 hr challenge with either TIGR4 or 6B. Total RNA was harvested and RT-PCR completed for 41 inflammatory genes. Heat mapped expression of A) 6B (MOI 20) challenged cells ± GSK-J4 (n=5), or heat mapped expression of B) TIGR4 (MOI 20) challenged cells ± IL-11 (n=4) to their respected uninfected condition. Multiple T-Test 6B to treatment and TIGR4 to treatment with significant genes highlighted in red. Principal component analysis of inflammatory RT-PCR panel using ΔCt comparing either C) TIGR4 (gray; n=5), 6B + 10 µM GSK-J4 (blue; n=5) and uninfected (n=5) or D) TIGR4 + 100 ng/mL IL-11 (gray; n=4), 6B (blue; n=5) and uninfected (n=5). Bioplot of the mean centered ΔCt data using the first two components and 60% concentration ellipses around each group.

In order to understand how KDM6B and IL-11 influenced generalized epithelial NF-κB responses, independently of pneumococcal-epithelial signaling events, we used IL-1β to stimulate A549 cells in the presence of either GSK-J4 or IL-11 (Sup. Fig. 4A). Inhibiting the catalytic activity of KDM6B lead to differential expression of multiple genes in comparison to untreated controls. In particular, there was a significant (pV≤0.015) 3-fold repression of the RelB transcript, a negative mediator of RelA induced pro-inflammatory responses ^47^, and a 0.6-fold increase in the IL-1β transcript, a potent pro-inflammatory cytokine. This comparison suggested inhibiting enzymatic activity of KDM6B lead to dysregulation of RelA/RelB cross talk and resolution. In contrast, exogenous IL-11 differentially regulated specific inflammatory mediators. We observed a significant reduction in *TNF* (pV≤0.030, 5-fold), another acutely pro-inflammatory cytokine. Thus, KDM6B and IL-11 are important regulators of cellular transcription in response to an inflammatory mediator IL-1β, as well as upon infection with pneumococci.

Finally, we used principal component analysis to compare how GSK-J4 or IL-11 treatment influenced the gene signatures induced by TIGR4 and 6B during infection. For GSK-J4, the ΔCt values from the gene panels of 6B + GSK-J4, TIGR4, and uninfected (untreated) were used for analysis. The first two components accounted for 53.8% of the variance observed between the treatment groups with three clusters forming across the first two components (60% confidence ellipses). Chemical inhibition of the catalytic activity of KDM6B resulted in the overlap of 6B + GSK-J4 treatment group with TIGR4 in the second dimension (Fig. 4C & Sup. Fig. 4B vector loadings; Sup. Table 3). Using the same approach, we compared the ΔCt values of TIGR4 + IL11, 6B, and uninfected (untreated). Here, the first two components explained 50.5% of the variation with three clusters of the experimental groups (60% confidence ellipses). Exogenous addition of IL-11 during TIGR4 challenge grouped and partially overlapped with the uninfected (untreated) and 6B groups (Fig. 4D & Sup. Fig. 4C vector loadings; Sup. Table 3). Altogether, these data suggest KDM6B and IL-11 play critical roles in moderating host transcription, particularly for a subset of NF-κB responses, and could potentially be used to exogenously interchange the host response to 6B and TIGR4 strains during pneumococcal infection.

### KDM6B and IL-11 contribute to epithelial cell integrity

Our RT-PCR expression data showed inhibition of KDM6B with GSK-J4 and exogenous IL-11 to moderate the host response. Interestingly, previous works demonstrate KDM6B and p65 are both required for keratinocyte wound healing ^22^. Using these findings, we hypothesized that KDM6B was involved with maintaining epithelial cell integrity during pneumococcal infection. To test the plasma membrane integrity of epithelial cells challenged with 6B or TIGR4 we used a Trypan blue exclusion assay, as damaged membranes are permissible to dye accumulation, in the presence of GSK-J4 or DMSO 24 hrs prior to challenge (Fig. 5A & B). We observed no difference in either epithelial integrity or cell viability between uninfected cells with GSK-J4 inhibitor by lactate dehydrogenase release (LDH; Fig. 5B & Sup. Fig. 5A). Furthermore, epithelial integrity was not compromised during 6B challenge in comparison to uninfected cells (Fig. 5A & B). In contrast, challenge with TIGR4 resulted in 60% epithelial membrane damage (Fig. 5B). Strikingly, inhibition of KDM6B catalytic activity affected epithelial integrity of 6B challenged cells, as shown by a significant 20% (pV≤0.001) increase in plasma membrane permissibility to Trypan entry (Fig. 5B). Since Pneumolysin, a pore forming cholesterol dependent cytolysin (CDC), is a well-documented and conserved virulence factor across all pneumococcal isolates and known to stimulate host responses, we tested if epithelial integrity loss was attributed to differential pneumolysin activity or expression between 6B and TIGR4 ^3,48^. We observed no difference in the hemolytic activity in the presence of GSK-J4 inhibitor, nor expression of pneumolysin between 6B and TIGR4 (Sup. Fig. 5B & C). Importantly, LDH assay showed not all Trypan blue positive cells were dead (Sup. Fig. 5A), which is in line with previous toxin studies ^49,50^. Overall, these results suggest KDM6B plays a role in mediating cell integrity during 6B challenge.

**Figure 5:**
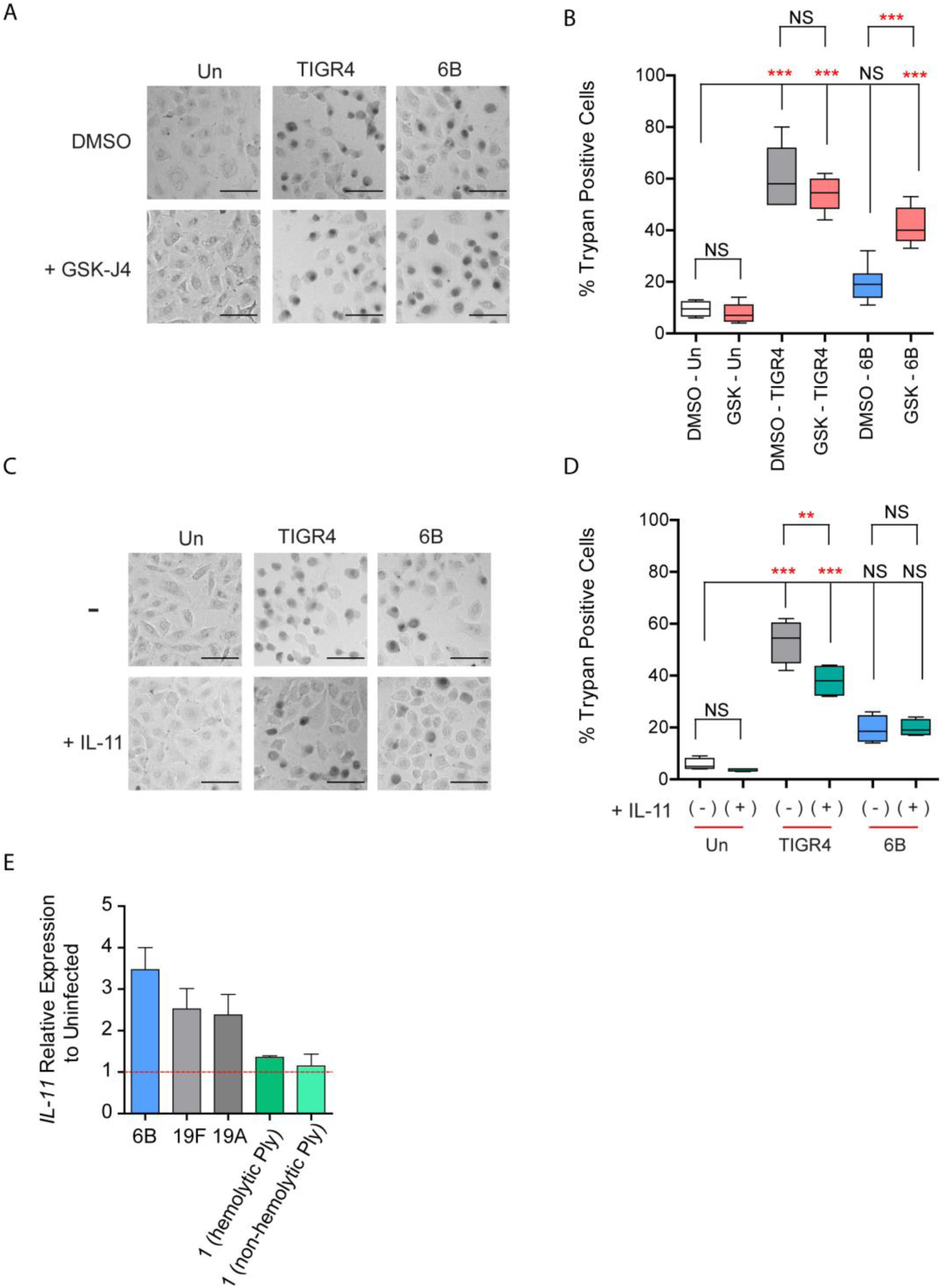
KDM6B and IL-11 contribute to epithelial cell integrity in response to pneumococcus. A549 cells untreated or treated with 10 µM GSK-J4 or 100 ng/mL recombinant human IL-11 prior to 2hr challenge with either TIGR4 (MOI 20) or 6B (MOI 20). Post-challenge cells were incubated with Trypan blue for cell integrity, fixed with 2.5% paraformaldehyde and imaged with a brightfield microscope. A) Representative image of GSK-J4 treated A549 cells post 2 hr challenge. Scale = 100µM. B) % Trypan positive cells between untreated and GSK-J4 treated (pink) (n=4; 200-300 cells per replicate and group). Uninfected white, TIGR4 gray and 6B blue. Tukey box and whisker plot. C) Representative image of IL-11 treated A549 cells post 2 hr challenge. Scale = 100 µM. D) % Trypan positive cells between untreated and IL-11 treated (green) (n=4; 200-300 cells per replicate and group). Uninfected white, TIGR4 gray and 6B blue. Tukey box and whisker plot. E) Total RNA harvested from A549 cells 2 hrs post-challenge with isolates of serotypes 6B, 19F, 19A, and two isolates of 1 (hemolytic and non-hemolytic Ply; n=3 per isolate). Relative expression of *IL-11* to uninfected cells. All data analyzed by One way-ANOVA with Tukey’s multiple comparison post-hoc test, **= pV≤0.01, ***= pV≤0.001, ns=not significant.

It is known respiratory tract epithelial cells can produce IL-11^51-53^, and IL-11 regulates epithelial cell plasma membrane proteins ^54^ while also modulating inflammatory, healing and mucosa responses ^55-58^. We therefore tested if exogenous addition of the cytokine IL-11 could mitigate the loss of epithelial cell integrity during TIGR4 challenge. At the time of challenge, the inoculums of TIGR4 and 6B were supplemented with recombinant human IL-11 cytokine prior to addition to A549 cells. After 2 hrs Trypan blue exclusion assay was performed (Fig. 5C & D). IL-11 at the time of TIGR4 challenge increased cell integrity by 20% (pV≤0.001), in comparison to untreated controls (Fig. 5D). There was no significant effect of exogenous IL-11 on uninfected or 6B challenged cells (Fig. 5D). Furthermore, LDH release showed exogenous IL-11 lowered TIGR4 cytotoxicity by 15% (pV≤0.01; Sup. Fig. 5A). Together this data shows IL-11 contributes to maintaining epithelial cell integrity during pneumococcal challenge.

Our data suggested the isolate of serotype 6B induced KDM6B and IL-11 to maintain epithelial integrity. We hypothesized other pneumococcal isolates commonly carried could also induce *IL-11* expression, while isolates associated with symptomatic disease and uncommonly carried would not. To test this, we compared isolates of serotypes 19A (Centre National de Référence des Pneumocoques) and 19F ^59^ (BHN100), which are associated with carriage, and two serotype 1 isolates harboring either a hemolytic or non-hemolytic pneumolysin allele, which are less commonly carried but cause disease states. Isolates of serotype 19A and 19F upregulated *IL-11* expression in A549 epithelial cells in comparison to uninfected cells, whereas the serotype 1 isolates did not (Fig. 5E). To determine if *IL-11* upregulation was specific to pneumococcal isolates, or potentially upregulated by other commensals, we tested five additional oral microbiome constituents. Indeed, *Streptococcus gordonii, Streptococcus sanguinis, Streptococcus oralis, Eikenella corrodens* and *Fusobacterium nucleatum* also upregulated the expression of *IL-11* in immortalized gingival keratinocytes (Sup. Fig. 5D). Together, our data show commonly carried pneumococcal isolates and individual commensal organisms induce *IL-11* expression, suggesting these bacteria induce a common response.

### KDM6B and IL-11 are essential for localized containment of pneumococcus *in vivo*

We established *in vitro*, inhibiting KDM6B and exogenous addition of IL-11 were sufficient to interchange cellular integrity to 6B and TIGR4. With this, we hypothesized local inhibition of KDM6B during 6B infection of the murine nasal epithelium would promote 6B to escape from the nasopharynx. We challenged mice with 3 - 4×10^6^ CFU of either 6B or TIGR4 mixed with either DMSO (vehicle control) or 5 mM GSK-J4. The plated inoculums showed no significant effect of DMSO or GSK-J4 on bacterial viability (Sup. Fig. 6B). Bacterial burden in the nasal lavage (NL), bronchoalveolar lavage fluid (BALF), lungs, and spleens of mice 24 hrs (Sup. Fig. 6A) and 48 hrs (Fig. 6A) post-inoculation were quantified by conventional colony forming unit (CFU) enumeration. Bacterial burdens from 6B and TIGR4 challenged DMSO animals showed on average one log more bacteria across all organs in comparison to TIGR4 by 24 hrs (Sup. Fig. 6A; 6B light blue; TIGR4 gray). By 48 hrs, infection with TIGR4 showed a progression of bacteria towards internal organs. The loosely attached bacteria in the NL and BALF decreased, while the burden in the lung and spleen increased in comparison to 24 hrs. However, 6B CFU numbers remained either constant or decreased in all samples (Fig. 6B; 6B light blue). With this data, we concluded 6B was primarily contained within the murine nasal cavity, whereas by 48 hrs post-inoculation TIGR4 had escaped the upper respiratory tract and begun to disseminate from the lungs.

**Figure 6:**
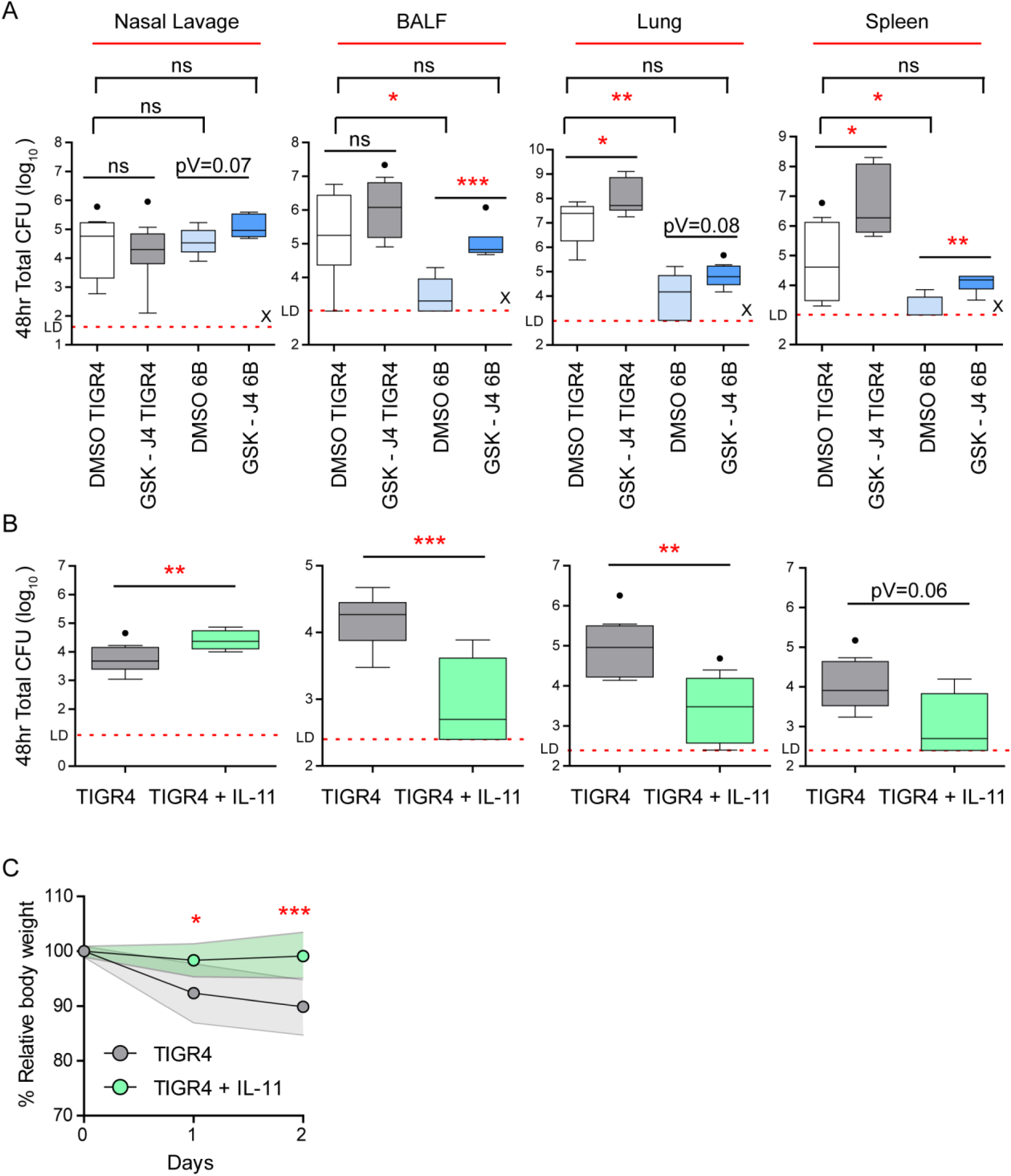
KDM6B is required for host response to serotype 6B. C57B6 mice (8-9 weeks) were challenged intranasally with 3 – 4×10^6^ CFU of TIGR4 or 6B supplemented with either DMSO (vehicle control), 5 mM GSK-J4 or 0.15 ug recombinant mouse IL-11. At indicated endpoints bacterial load was enumerated by conventional CFU counts on 5 µg/mL Gentamicin Columbia Blood agar selection plates from the nasal lavage (NL), bronchoalveolar lavage fluid (BALF), lungs, and spleen of infected animals. A) 6B and TIGR4 ± GSK-J4 with matched DMSO controls challenged animals 48 hrs post-inoculation CFU burden of indicated samples (n=9). DMSO TIGR4 white, GSK-J4 TIGR4 gray, DMSO 6B light blue, GSK-J4 6B dark blue. B) TIGR4 ± IL-11 challenged animals 48 hrs post-inoculation CFU burden of indicated samples (n=9). TIGR4 (gray; n=9) and TIGR4 + IL-11 (green; n=9). Tukey box and whisker plot with dots representing outliers. One way-ANOVA non-parametric Kruskal-Wallis with Dunn’s multiple comparison post-hoc test, *= pV≤0.05, **= pV≤0.01, ***= pV≤0.001, ns=not significant. CFU=colony forming unit. Dotted lines =Limit of detection (LD). LD for each organ: NL (50 CFU); BALF, Lung and Spleen (1000 CFU). X indicates animal died prior to endpoint. C) Percent initial body weight TIGR4 gray and TIGR4 + IL-11 green. Solid lines represent Std. Dashed line indicates 20% weight loss threshold. Repeated measures ANOVA with Tukey’s multiple comparison post-hoc test, **= pV≤0.01, ***= pV≤0.001.

However, the addition of GSK-J4 changed the bacterial distribution of 6B. Indeed, GSK-J4 treated animals challenged with 6B showed increased burden across all samples in comparison to the 6B DMSO control group at 24 hrs (Sup. Fig. 6A). Additionally, the recovered bacteria from the NL and BALF in the GSK-J4 6B challenged group was not significantly different from the TIGR4 DMSO group (Sup. Fig. 6A). However, after 24 hrs there was no significant difference in the recovery of TIGR4 from the NL, BALF, lungs or spleens between DMSO or GSK-J4 treated animals (Sup. Fig. 6A; TIGR4 DMSO white, TIGR4 GSK-J4 gray). By 48 hrs post-challenge 6B GSK-J4 animals maintained a significantly (pV≤0.05) high bacterial burden in the BALF and spleen compared to 6B DMSO treated animals (Fig. 6A; 6B DMSO light blue, 6B GSK-J4 dark blue). Importantly in animals treated with GSK-J4 6B, was recovered from the spleen, an organ in which bacteria were mostly undetected in DMSO control animals. GSK-J4 treated animals in the TIGR4 group also showed an increase in bacterial burden at 48 hrs post-challenge in the NL, BALF, lung and spleen, which suggests a basal role of KDM6B in regulating NF-κB responses. Altogether, these data show KDM6B activity is specifically required for localized containment of 6B during infection of the murine nasal cavity, and is potentially a negative regulator of TIGR4 dissemination.

In our previous *in vitro* studies exogenous IL-11, at the time of infection, was sufficient to modify the host response to TIGR4. This suggested that localized IL-11 during TIGR4 challenge *in vivo* could potentially delay translocation of the bacteria into the lower respiratory tract and deeper tissues by promoting localized containment in the upper respiratory tract. To address this, animals were intranasally challenged with 3 – 4×10^6^ CFU of TIGR4 with or without recombinant mouse IL-11 (final 0.15 ug per mouse; Sup. Fig. 6C) at the time of infection. Supplementation of midlog phase TIGR4 inoculums had no effect on growth (Sup. Fig. 6D). Animal weight, as a read out of symptomatic disease, was monitored over 48 hrs, and the bacterial burden enumerated 48 hrs post-infection from the NL, BALF, lung and spleen (Fig. 6B). By 48 hrs post-infection there was substantial recovery of TIGR4 bacterial burden in the BALF, lung and spleen (Fig. 6B), which accompanied a 10% loss of initial starting weight (Fig. 6C). Together this was indicative of symptomatic pneumococcal disease induced by TIGR4 infection. In contrast, TIGR4 with recombinant mouse IL-11, at the time of intranasal challenge, resulted in 5-fold retention of bacteria within the NL (Fig. 6B; pV≤0.01) in comparison to TIGR4 infection alone. Dissemination of bacteria to internal organs was also diminished, with greater than a one-log difference on average in bacterial burden across the BALF, lung and spleen (Fig. 6B). These bacterial burdens further coincided with a significant 5-10% preservation of initial starting weight at 24 hrs and 48 hrs post-infection in contrast to the TIGR4 group (Fig. 6C; pV≤0.05 and pV≤0.001 respectively). Overall, these data demonstrate during TIGR4 pneumococcal infection a localized IL-11 response is sufficient to ameliorate symptomatic disease and reduce bacterial dissemination to the lower respiratory tract.

## Discussion

Bacterial contact with the epithelial barrier is an essential process that leads to pneumococcal carriage and precedes symptomatic pneumococcal disease ^3,5^. However, the molecular and transcriptional processes across the spectrum of host interaction from nasal microbiome constituent to potentially lethal disease states are not completely understood. Towards this end, we first demonstrated the pneumococcal strain 6B ST90 (CC156 lineage F) did not cause symptomatic disease or lethality in contrast to a serotype 4 (TIGR4) strain in a murine model of infection. Using these two strains, which induce opposing host outcomes, we completed a human microarray of A549 epithelial cells challenged with either 6B ST90 or TIGR4 pneumococcal strains to better understand the cellular processes involved. We show a serotype 6B isolate differentially regulated 388 genes, with enrichment for NF-κB associated genes in comparison to a TIGR4 strain. Direct comparison of NF-κB regulated genes between 6B and TIGR4 showed that 6B induced a unique transcriptional response that included *KDM6B* and *IL-11* expression, which required phosphorylation of p65. We demonstrate molecularly that the 6B isolate, through the activity of KDM6B, induces remodeling of the IL-11 promoter to reveal three NF-κB sites, which are not accessible during IL-1β or TIGR4 stimulation. Together, this is the first demonstration that a 6B strain remodels chromatin within epithelial cells to support containment in the upper respiratory tract.

The lysine demethylase KDM6B has mainly been characterized in cellular development, however a few studies suggest that this particular histone demethylase also fine-tunes inflammatory responses and wound healing downstream of p65 through largely unknown mechanisms ^20-23,60-62^. We are the first to report both a biological and molecular role for KDM6B and H3K27me3/2 in regulation of a specific gene locus, IL-11, during bacterial challenge. Surprisingly, although KDM6B was shown to primarily target H3K27me3 and to a lesser extent H3K27me2 ^37,38,46^, our results suggest KDM6B is selectively demethlyating H3K27me2 and not H3K27me3 at the IL-11 promoter. These results are consistent with previous observations of *Da Santa et al.*, who reported gene regulation by KDM6B independently of H3K27me3 ^20^. With this observation, we hypothesize that KDM6B is differentially regulating inflammatory gene expression by selective demethylation of H3K27me2 through either an unknown regulatory element or posttranslational modifications to KDM6B. Future ChIP-seq and proteomic studies with biological stimuli, such as pneumococcus, will yield substantial insight into possible KDM6B complexes, and the dynamics of H3K27me3/2 in epigenetic control of inflammatory signaling cascades.

Our findings support the idea that pneumococcal-epithelial interactions may actively induce host responses that promote confinement of bacteria to the upper respiratory tract, leading eventually to clearance. The host inflammatory response is a known contributor to both the commensal and pathogenic lifestyles of pneumococcus, by playing an important role in clearance, transmission and establishment of a replicative niche ^3,13,63^. Our results suggest KDM6B, and its regulation of *IL-11* transcription, are key components modulating a localized epithelial host response. In a murine model, chemical inhibition of KDM6B led to the dissemination of an otherwise contained 6B strain to the spleen. Given the role of KDM6B, we show that containment is a host driven mechanism involving the expression of IL-11. Since KDM6B differentially regulates multiple genes, we cannot rule out the possibility there are others with concurrent or synergistic functions with IL-11. However, our IL-11 rescue experiments with TIGR4 suggest a role for IL-11 in locally maintaining a host-limiting/tolerogenic epithelial-pneumococcal host response. Interestingly, IL-11 is known to influence mucus production, cell membrane components, wound healing of gastric ulcers and resistance of endothelial cells to immune mediated injury ^54,56,57,64,65^. Promoting this localized response to our serotype 6B ST90 isolate within the nasopharynx, is also reflected in our microarray data, as 52 genes associated with wound healing gene ontology were upregulated by 6B in comparison to TIGR4 (Sup. Table 1). Additionally, 39 of the 52 upregulated wound healing genes were also associated with KDM6B as determined by ChIP-seq studies of macrophages (Sup. Table 1). Within this principle, we propose that some pneumococcal strains, including the 6B strain tested, along with some bacteria of the oral microflora, actively influence their localization within the host through a subset of NF-κB driven “wound healing” genes as a potential means to counter balance a deleterious pro-inflammatory host response within the host. How the localized inflammatory processes and host-mediated confinement influences carriage duration and clearance, transmission, or the impact of micro invasion ^13^ upon these processes remains to be defined.

Through our study of p65 activation by pneumococcus, we find that the 6B isolate activated p65 to similar levels as IL-1β, however, the ensuing transcriptomic responses are different. Combining these observations with active remodeling of the IL-11 promoter suggests that under 6B stimulation there are additional p65 interacting partners or posttranslational modifications (PTMs), in conjunction with phosphorylation of serine 536. Such data would support a biological role for “NF-κB barcode hypothesis”, where a signature barcode of PTMs on NF-κB mediates a specific gene expression pattern ^66,67^. While we have established a link between p65, KDM6B and *IL-11* expression, identification of the p65 PTMs and interacting partners necessary for the NF-κB dependent processes of 6B will advance our understanding of not only pneumococcal induced host responses, but also p65 regulation during tolerogenic inflammatory responses.

A meta-analysis conducted by *Brouwer et al*., highlighted single nucleotide polymorphisms (SNPs) associated with *NFKBIA, NFKBIE*, and *TIRAP* correlated with protection, whereas SNPs within NEMO (*IKBKG)* or IRAK4 associated predominantly with increased susceptibility to disease ^68^. Analysis of KDM6B and H3K27me3 ChIP-seq data from LPS stimulated macrophages, shows these protective genes, *NFKBIA, NFKBIE*, and *TIRAP*, are also associated with KDM6B and/or H3K27me3, whilst NEMO and IRAK4 are not ^20^. Since *NFKBIA*, and *NFKBIE* are known to inhibit NF-κB through sequestration within the cytoplasm ^15,69,70^, one could hypothesize KDM6B is a chromatin level negative regulator that balances inflammatory signaling in conjunction with p65 across a unique “p65-KDM6B” axis. In this context, KDM6B serves as the molecular “regulator or brake” responsible for modulating the host response based upon the severity, or degree of inflammatory signal input. Interestingly, we found when IL-1β was used to stimulate epithelial cells pretreated with GSK-J4, to inhibit KDM6B activity, there was a significant loss of the *RelB* transcript, a known negative mediator of RelA ^47^. How KDM6B influences *RelB* transcription is unknown. Further support for KDM6B as an inflammatory regulator is evidenced by our *in vivo* studies showing chemical inhibition of KDM6B dampens the ability of the murine host to control TIGR4 infection and the escape of an otherwise confined serotype 6B isolate from the murine nasal cavity.

Overall, our data demonstrates the first biological role of KDM6B in moderating the host response to bacteria. We further reveal catalytically active KDM6B is required for host-mediated tolerogenic confinement of a 6B ST90 (CC156 lineage F) strain within a murine model of infection. We further show exogenous addition of IL-11, in the same murine model, to promote confinement of an otherwise lethal infection with a TIGR4 strain. Future studies characterizing the molecular interplay of chromatin and cellular processes across the spectrum of pneumococcal host responses will not only identify new means to combat pneumococcal disease, but may also reveal the processes exploited by commensal organisms as well.

## Materials and Methods

### Bacterial strains, growth conditions and CFU enumeration

Clinical isolates of serotypes 6B (ST90; CNRP# 43494), 19A (ST276; CNRP# 45426) and 1 (non-hemolytic; ST306; CNRP# 43810) were obtained from the Centre National de Référence des Pneumocoques (Emmanuelle Varon; Paris, France). Serotype 19F (BHN100; ST162 Birgitta Henriques Normark, Karolinska Institutet ^59^), serotype 4 TIGR4 (Thomas Kohler, Universität Greifswald), and serotype 1 (ST304 hemolytic; M. Mustapha Si-Tahar, Université de Tours). Experimental starters were prepared from frozen master stocks struck on 5% Columbia blood agar plates (Biomerieux Ref# 43041) and grown overnight at 37°C with 5% CO_2_ prior to outgrowth in Todd-Hewitt (BD) broth supplemented with 50 mM HEPES (Sigma) (TH+H) at 37°C with 5% CO_2_ in closed falcon tubes. Midlog bacteria were pelleted, and diluted to 0.6 OD_600_ /mL in TH+H media supplemented with Luria-Bertani (BD) and 15% glycerol final concentration. Aliquots were made and frozen at −80°C for experiments. All experiments were performed with frozen experimental starters of *S. pneumoniae* less than 14 days old. For experiments, starters were grown to midlog phase in TH+H broth at 37°C with 5% CO_2_ in closed falcon tubes, pelleted at 1,500xg for 10 mins at room temperature (RT), washed in DPBS, and concentrated in 1mL DPBS prior to dilution at desired CFU/mL using 0.6 OD_600_ /mL conversion factors in either cell culture media or DPBS for animal studies (conversion factors Sup. Table 4). For paraformaldehyde (PFA) killed bacteria the concentrated bacteria, prior to dilution, was incubated 4% PFA for 30 mins at RT, washed in DPBS, and diluted to desired CFU/mL using 0.6 OD_600_ /mL conversion factors. Bacteria CFU enumeration was determined by serial dilution plating on 5% Columbia blood agar plates.

### Cell culture and *In vitro* challenge

A549 human epithelial cells (ATCC ref# CCL-185) were maintained in F12K media (Gibco) supplemented with 1x GlutaMax (Gibco) and 10% heat inactivated fetal calf serum (FCS) at 37°C with 5% CO_2_. Detroit 562 nasopharyngeal epithelial cells (ATCC ref# CCL-138) and BEAS-2B epithelial cells (ATCC ref# CRL-9609) were maintained in DMEM supplemented with 1x sodium pyruvate (Gibco) and 1x GlutaMax (Gibco) 10% heat inactivated FCS. Stable HeLa GFP-p65 were generated using the transposon-based sleeping beauty system, and maintained in DMEM supplemented with 1x sodium pyruvate (Gibco), 1x GlutaMax (Gibco) and 10% heat inactivated FCS ^71^. A549, BEAS-2B, Detroit 562 or HeLa GFP-p65 cells were used until passage 15. For *in vitro* challenge studies, A549, BEAS-2B, Detroit 562, or HeLa GFP-p65 cells were plated in tissue culture treated plates at 2×10^5^ cells (6well; for 72 hrs), 5×10^4^ cells (24well; for 48 hrs), or 1×10^4^ cells (96well; for 48 hrs). Cells were used at 90% confluency for all *in vitro* studies. Bacterial inoculums diluted in cell culture media were added to cells, and bacterial-epithelial cell contact synchronized by centrifugation at 200xg for 10 mins at RT, then moved to 37°C with 5% CO_2_ for 2 hrs. For inhibitor studies, cell culture media was aspirated, and replaced with filter sterilized culture media containing inhibitor volume matched DMSO (Sigma), GSK-J4 (Sigma ref# SML0701), or BAY 11-7082 (Sigma ref# B5556) at 10 µM final concentration for 24 hrs or 3 hrs respectively prior to bacterial addition. Human IL-11 (Miltenyi Biotec ref# 130-094-623) and human IL-1β (Enzo Life Sciences ref# ALX-522-056) were used at 100 ng/mL and 10 ng/mL final concentration respectively in cell culture media.

### Immunofluorescence and Trypan blue bright field microscopy

To quantify nuclear KDM6B, A549 cells were seeded on acid washed and UV treated coverslips in 24well plates as described above, 2 hrs post-challenge media was aspirated, cells washed in DPBS, and fixed with 2.5% PFA for 10 mins at RT. Coverslips were blocked and permeabilized overnight in 5% BSA 0.5% Tween20. Coverslips were incubated for 1hr at RT with KDM6B (1:500; abcam ref# ab38113) diluted in 5% BSA 0.5% Tween20, washed three times in 0.5% Tween20, and incubated for 1 hr with Alexa Fluor 488 secondary. After secondary, coverslips were washed three times in 0.5% Tween20 and mounted using Prolong Gold with DAPI (Invitrogen). Confocal microscopy images were acquired on a Zeiss axio observer spinning disk confocal. Nuclear KDM6B intensity per cell was quantified within an ROI generated from the DAPI signal in Fiji ^72^. For Trypan exclusion microscopy, A549 cells were seeded in 96well plates as described above. 2 hrs post-challenge culture media was aspirated, cells washed in DPBS, and Trypan blue (Thermo) added for 10 mins at RT. Trypan blue was removed, and cells fixed with 2.5% PFA for 10mins at RT. PFA was removed and fixed cells washed in DPBS prior to imaging on a EVOS FL (Thermo). Trypan positive cells were scored manually as % of total cells in an imaged field.

### A549 epithelial microarray

A549 cells were infected as described above, and total RNA harvested using RNeasy kit (Qiagen). RNA quality was confirmed using a Bioanalyzer (Agilent). Affymetrix GeneChip human transcriptome array 2.0 was processed as per manufacturer’s instructions. Data was analyzed using TAC 4.0 (Applied Biosystems).

### LDH assay

LDH assays were performed on cell culture supernatants as per manufacturer’s instructions (Pierce LDH cytotoxicity kit ref# 88953). LDH absorbance was read using Cytation 5 (BioTek) at manufacturer’s recommended excitation and emissions.

### ChIP and ChIP-qPCR

Detailed ChIP buffer components are in supplemental methods. In brief, 8×10^6^ A549 cells were cross-linked in tissue culture plates with 1% formaldehyde for 10 mins at RT, then quenched with 130 mM glycine for 5 mins at RT. Cells were washed in DPBS, gently scraped, and transferred to an eppendorf. Harvested cells were pelleted at 200xg, supernatant aspirated and frozen −20°C. To obtain chromatin, cell pellets were thawed on ice and lysed for 30 mins on ice in nuclear isolation buffer supplemented with 0.2% Triton X-100. Nuclei pelleted, supernatant aspirated and suspended in chromatin shearing buffer for sonication (10 cycles of 15 sec ‘on’ and 30 sec ‘off’) with a Bioruptor (Diagenode) to 200-900bp size. Sheared chromatin was cleared by centrifugation, then sampled for size using 2% agarose gel electrophoresis and quantification using Pico488 (Lumiprobe ref# 42010). ChIP grade antibodies to p65 (L8F6) (CST ref #6956), KDM6B (abcam ref# ab38113), H3K27me3 (abcam ref# ab6002), H3 (abcam ref# ab195277), or H3K27me2 (diagenode ref# C15410046-10) were used at manufacturer’s recommended concentrations and bound to DiaMag beads (diagenode ref # C03010021-150) overnight with gentle rotation. Quantified chromatin was diluted to 10 µg per immunoprecipitation condition was added to antibody bound DiaMag beads overnight with gentle rotation with 8% of input reserved. Beads were washed with buffers 1-6 (supplemental methods), decrosslinked by boiling for 10 mins with 10% Chelex, treated with RNase and proteinase K, then purified using phenol-chloroform extraction followed by isopropanol precipitation. Recovered DNA was suspend in molecular grade water, and 1 µL used for Sybr Green reactions as per manufacturer’s instructions on a BioRad CFX384 (BioRad). % recovery was calculated as 2 raised to the adjusted input Ct minus IP Ct multiplied by 100. For histone marks, H3K27me3/2, the % recovery was normalized to the % recovery of H3. ChIP qPCR primers listed in Sup. Table 4.

### RNA isolation and RT-PCR

Total RNA isolated using TRIzol (Life technologies ref#15596-026) extraction method as per manufacturer’s recommendations. Recovered RNA was suspended in molecular grade water, nano dropped and 5 µg converted to cDNA using Super Script IV as per manufacturer’s instructions. cDNA was diluted to 20 ng/µL in molecular grade water and 1 µL used for Sybr Green reactions in technical duplicate or triplicate as per manufacturer’s instructions on a BioRad CFX384 (BioRad). RT-PCR primers listed in Sup. Table 4. Relative expression was calculated by ΔΔCt method to *GapDH* ^73^.

### Immunoblots and quantification

Cell culture media was removed, cells were washed in DPBS and whole cell lysates harvested with Laemmli buffer ^74^. Lysates were boiled 10min, and frozen at −20°C. Whole cell lysates were ran on 8% polyacrylamide SDS PAGE gels, transferred to PVDF membrane (BioRad TransBlot), blocked 1 hr in 5% BSA TBST, then probed for p65 (CST ref #6956), p65 phosphorylation at serine 536 (CST ref# 3033), or actin AC-15 monoclonal (Sigma ref# A5441) as per manufacturer’s recommendations. Appropriate secondary-HRP conjugated antibodies were used with clarity ECL (BioRad) developing reagents. Membranes were developed on ChemiDoc Touch (BioRad).

### *In vivo* animal studies

All protocols for animal experiments were reviewed and approved by the CETEA (Comité d’Ethique pour l’Expérimentation Animale - Ethics Committee for Animal Experimentation) of the Institut Pasteur under approval number Dap170005 and were performed in accordance with national laws and institutional guidelines for animal care and use. Wildtype C57BL/6 female 8-9 week old mice were purchased from Janvier Labs (France). Wildtype C57BL/6 male 10-12 week old mice used for survival studies (Institut Pasteur). Animals were anesthetized with a ketamine and xylazine cocktail prior to intranasal challenge with 20 µL of 3 – 4×10^6^ CFU of TIGR4 or 6B. Bacterial inoculums were made as described above with minor modification. In brief, after 0.6 OD_600_ bacterial were concentrated and diluted in either filter sterilized DMSO/DPBS or 5 mM GSK-J4/DPBS. For IL-11 studies, recombinant mouse IL-11 (Enzo; ENZ-PRT243-0010) was added to bacterial inoculums to a final concentration of 7,500 ng/mL (0.15 ug per mouse). After 24 hrs or 48 hrs post-inoculation animals were euthanized by CO_2_ affixation. The nasal lavage was obtained by blocking the oropharynx, to avoid leakage into the oral cavity and lower airway, and nares flushed with 500 µL DPBS. Bronchoalveolar lavage fluid (BALF), lungs, and spleens were collected and placed in 1 mL DPBS supplemented with 2x protease inhibitor cocktail (Sigma ref # P1860). CFU were enumerated as described above on 5 µg/mL Gentamicin Columbia Blood agar selection plates. For survival studies, animals were monitored daily for weight loss and disease signs. Animals were euthanized by CO_2_ affixation at ≥20% loss of initial weight or at body conditioning score (BCS) ≤2^75^.

### Statistical analysis

All experiments, unless otherwise noted, were biologically repeated 3-4 times with the statistical test in the figure legends. Data normality was tested by Shapiro-Wilk test, and appropriate parametric or non-parametric tests performed. *P* values were calculated using GraphPad Prism software. For RT-PCR all statistics were calculated on either the ΔΔCt or ΔCt depending on the desired comparison. PCA plots using the “prcomp” function of the base stats package in R on scaled and mean centered log_2_ transformed data, or ΔCt for cross-treatment comparisons. Microscopy data was collected from analysis of 30-100 cells for nuclear staining, or 200-300 cells for brightfield per biological replicate per group. ChIP-qPCR studies used four technical replicates per biological with all data graphed. Animal studies used the minimum number of animals required for power calculated by analysis of CFU using G*Power software.

## Supplemental Methods

### Human IL-11 ELISA

Detroit 562 cells were infected with *S. pneumoniae* for 6 hrs, and supernatants were collected for IL-11 analysis using an IL-11 Human Elisa Duo Set (R&D Systems, DY218 lot number 5225707), according to manufacturers’ instructions.

### AlamarBlue cytotoxicity

A549 cell viability was determined using AlamarBlue (Thermo ref# DAL1025) as previously described ^76^. AlamarBlue absorbance was read using a Cytation 5 (BioTek) at manufacturer’s recommended excitation and emissions.

### Crude hemolytic assay

Red blood cells (RBC) from horse blood (Thermo ref# SR0050C) were washed three times in DPBS and suspended at a final concentration of 1% v/v, with 100 µl moved to individual wells of a 96well U-bottom plate and stored at 4°C until use. Cell culture supernatants were collected from infected A549 cells 2 hrs post-infection, centrifuged and filtered prior to 100 µl being combined with RBCs. For positive lysis control, volume matched water and 2% triton-X100 were used, the negative control was RBCs with cell culture media alone. The plate was incubated for 30 mins at 37 °C with 5% CO_2_, then centrifuged 5 mins at 200xg. Supernatants (100 µl) were transferred to a new clear 96well plate and absorbance (OD_540_) read using Cytation 5 (BioTek). % Lysis was calculated as absorbance of sample adjusted for media alone over positive lysis sample adjusted for media alone multiplied by 100.

### Oral commensal *in vitro* infection and RT-PCR

*Streptococcus gordonii, S. oralis*, and *S. sanguinis* were all cultivated in brain heart infusion (BHI) broth, *Fusobacterium nucleatum* was grown in BHI broth supplemented with 5 ug/mL hemin and 1 ug/mL menadione, and *Eikenella corrodens* was grown in TYGVS broth. All bacteria were cultured anaerobically until midlog phase. Human telomerase immortalized gingival keratinocytes (TIGK) were maintained at 37 °C with 5% CO_2_ in Dermalife-K serum-free culture medium (Lifeline Cell Technology)^77^. Epithelial cells at 80% confluency were challenged with the different bacteria species at an MOI of 100 for 3 hrs, were washed and fresh media was added for another 18 hrs. Total RNA was isolated using the Qiagen RNeasy mini kit (Qiagen). RNA concentrations were determined spectrophotometrically using a NanoDrop 2000 and quality was determined using the Qubit 4 fluorimeter. cDNA from total RNA was synthesized (500 ng RNA per reaction volume) using a High Capacity cDNA reverse transcription kit (Applied Biosystems). qRT-PCR was performed using *IL-11* and *GAPDH* primers in Sup. Table 4. qRT-PCR was performed on an Applied Biosystems QuantStudio 3 cycler and the auto-calculated threshold cycle selected. The cycle threshold (Ct) values were determined, and mRNA expression levels were normalized to *GAPDH* and expressed relative to controls following the 2-ΔΔCT method.

### Pneumococcus growth curve with recombinant murine IL-11

Midlog bacterial inoculums for both TIGR4 and TIGR4 supplemented with IL-11, as described under “*In vivo* animal studies”, were used for growth analysis with the final dilution being in TH+H. Bacterial aliquots (100µl each) were distributed into wells of a clear 96well plate. Plates were incubated at 37°C with 5% CO_2_ for 2 hrs, and absorbance (OD_600_) taken at 20 min intervals using a Cytation5 (BioTek). Absorbance was calculated after normalizing to TH+H growth media alone.

ChIP buffer solutions as follows:

| Nuclear Isolation Buffer (NIB) | Chromatin shearing buffer (Buffer C) | SDS dilution buffer (buffer D) |
| --- | --- | --- |
| 15mM Tris pH 7.5 | 1% SDS | 0.6% Triton X-100 |
| 60mM KCl | 10mM Tris HCL pH 8.0 | 0.06% NaDOC |
| 15mM NaCl<sub>2</sub> | 1mM EDTA | 150mM NaCl |
| 250mM sucrose | 0.5mM EGTA | 12mM Tris HCL pH 8.0 |
| 1mM CaCl<sub>2</sub> |  | 1mM EDTA |
| 5mM MgCl<sub>2</sub> |  | 0.5mM EGTA |
| Day of use | Day of use | Day of use |
| 1x PhosSTOP | 1x PhosSTOP | 1x PhosSTOP |
| 10mM Sodium Butyrate | 10mM Sodium Butyrate | 10mM Sodium Butyrate |
| 0.2mM PMSF | 0.2mM PMSF | 0.2mM PMSF |
| 1x protease cocktail | 1x protease cocktail | 1x protease cocktail |
| 1x phosphatase inhibitor cocktail | 1x phosphatase inhibitor cocktail | 1x phosphatase inhibitor cocktail |

| 10% Chelex | 1.25M pH 7.6 Glycine buffer |
| --- | --- |
| 10g into 100mL miliQ water | 46.91g Glycine in 400mL milliQ water |
|  | pH adjusted to 7.6 and QC to 500mL |

| Wash buffer 1 (isotonic) | Wash buffer 2 (isotonic, ionic charge change) | Wash buffer 3 (high salt dilution) |
| --- | --- | --- |
| 1% Triton X-100 | 0.5% NP40 | 0.7% Triton X-100 |
| 0.1% NaDOC | 0.5% Triton X-100 | 0.1% NaDOC |
| 150mM NaC | 0.5% NaDOC | 250mM NaCl |
| 10mM Tris HCL pH 8.0 | 150mM NaCl | 10mM Tris HCL pH 8.0 |
|  | 10mM Tris HCL pH 8.0 |  |
| Wash buffer 4 (high salt dilution) | Wash buffer 5 (salt dilution) | Wash buffer 6 (TE) |
| 0.5% NP40 | 0.1% NP40 | 20mM Tris HCL pH 8.0 |
| 0.5% Triton X-100 | 150mM NaCl | 1mM EDTA |
| 250mM LiCl | 20mM Tris HCL pH 8.0 |  |
| 20mM Tris HCL pH 8.0 | 1mM EDTA |  |
| 1mM EDTA |  |  |

## Supporting information

supplemental figures supporting main figures

## Author Contributions

Conceived and designed all experiments: MGC and MAH. Performed experiments: MGC; EP (KDM6B microscopy and HeLa GFP-p65 western blot); TC and OR (animal studies); CMW (IL-11 ELISA); DPM and RJL performed all oral commensal infections and RT-PCR. Analyzed data: MGC. LB generated the stable HeLa GFP-p65 cell line under the supervision of JE. MGC wrote the original manuscript draft. MGC and MAH edited and reviewed the manuscript. MAH supervised the research. All authors approved the final manuscript.

## Acknowledgments

We would like to thank Emmanuelle Varon, Birgitta Henriques Normark, M. Mustapha Si-Tahar, and Thomas Kohler for their generous gifts of *S. pneumonie* strains. We are thankfully Gregory Dore (Institut Pasteur) for processing the microarray Affymetrix GeneChips. Biostatistics and R discussions with Jose Nabuco (Institute Pasteur) and Jeremy Camp (University of Veterinary Medicine Vienna) were greatly appreciated. Michael G. Connor is supported by the Pasteur Foundation Fellowship. T.M.-N.C. is supported by a fellowship from the Foundation for Medical Research (Mariane Josso Prize), and a donation from Credit Agricole. Melanie Hamon, G5 Chromatin and infection, is supported by the Institut Pasteur, and the Agence National de la Recherche (ANREpiBActIn). Caroline M. Weight was supported by the Medical Research Council (MR/T016329/1). We would like to thank Robert S. Heyderman (UCL) for in depth discussion on the manuscript and is supported by the MRC (MR/T016329/1). Jost Enninga and Laura Barrio, members of Dynamics of host-pathogen interactions unit (Institut Pasteur), are supported by the European Commission (ERC-CoG-Endosubvert), the ANR-HBPsensing, and are members of the IBEID and Milieu Interieur LabExes. Richard J. Lamont is funded by NIH/NIDCR DE01111, DE012505, DE017921 and DE023193.

