## supplemental figures supporting main figures for "The histone demethylase KDM6B fine-tunes the host response to *Streptococcus pneumoniae*"

**Figure Legends and Tables:**

**Supplemental Figure 1:** *Human microarray, KDM6B microscopy and demethylase RT-PCR.* A) TIGR4 and 6B inoculums for animal intranasal challenge model (n=3). B) Human microarray of epithelial A549 cells 2 hrs post-challenge (MOI 20) with either 6B (blue) or TIGR4 (gray). All genes differentially regulated by ± 1.5 fold-change to uninfected condition. Circled 6B genes are NF-κB associated. C) Comparison of genes identified by microarray containing NF-κB sites between 6B and TIGR4. D) Clustering heat map, using ClustVis [1], (Euclidean for genes, Correlation for cell line) of the median $\log_2$ relative expression RT-PCR data for indicated genes from A549 (n=5), Detroit 562 (n=2) and Beas2b (n=1) cell lines. KDM6B and IL-11 cluster highlighted in red. E) Representative images of KDM6B (green) merged with nucleus stained with DAPI (false colored red for visual display). Scale = 10 µm. F) Quantified IL-11 ELISA (n=5; 4-6 replicates per biological) 6 hrs post-infection of Detroit 562 cells. Median centered (red) dot plot of all values. Dashed redline is the limit of detection. G) Total RNA 2 hrs post-challenge with 6B or TIGR4 from A549 cells. Demethylase panel RT-PCR shown as ΔCt (n=3; 3 technicals per biological replicate). Tukey box and whisker plot. Data in A) and G) analyzed by One way-ANOVA with Tukey's multiple comparison post-hoc test, **= pV≤0.01, ***= pV≤0.001, ns=not significant.

**Supplemental Figure 2:** *Inhibitor viability and Pneumolysin data.* A549 cells untreated or treated with 10 µM BAY 11-7082, 10 µM GSK-J4, or DMSO vehicle control (n=4). A) Cell viability determined by AlamarBlue. Expressed as % of untreated cells. One way-ANOVA with Dunnett's multiple comparison post-hoc test to untreated, no significant difference observed. B) Transcript levels for *IL-11*, *KDM6B* and *PTGS2* determined by RT-PCR. Displayed as ΔCt for comparison to untreated (n=4). Bar graph ± Std. All data analyzed by One way-ANOVA with Tukey's multiple comparison post-hoc test, no significant difference observed.

**Supplemental Figure 3:** *ChIP-PCR KDM6B locus and IL-11 locus for TIGR4 and IL-1β.* Chromatin obtained from untreated and 10 µM GSK-J4 treated A549 cells 2 hrs post-challenge with 6B (MOI 20), TIGR4 (MOI 20) or IL-1β. 10 µg chromatin input used for ChIP of p65, KDM6B, H3K27me3 and histone H3 (H3), followed by ChIP-qPCR at primer locations (P6, P3 & P2) spanning the NF-κB sites upstream of the transcriptional start site (TSS). A) Schematic of KDM6B promoter with ChIP-qPCR primer locations (P4 & P3) and the NF-κB sites. B) % recovery of input for p65 2 hrs post-infection with 6B (n=3 untreated; n=3 GSK-J4 treated). C) Schematic of IL-11 promoter with ChIP-qPCR primer locations (P6, P3 & P2) and the NF-κB sites. D - G) % recovery of input for p65, KDM6B, H3K27me3 normalized to H3, or H3 bound at P6, P3 & P2 in untreated and GSK-J4 treated samples (n=3 untreated; n=3 GSK-J4 treated). Tukey box and whisker plot with dots representing outliers. One way-ANOVA with Tukey's multiple comparison post-hoc test, no significant difference observed in comparison to uninfected/untreated or 6B samples from Fig. 3.

**Supplemental Figure 4:** *IL-1β RT-PCR with GSK-J4 or IL-11 treatments.* A) Heat map represents fold change to either untreated or treatment matched controls per condition (n=5 for untreated, and n=3 per treatment). Heat mapped difference between relative expression of treatments to untreated. Multiple T-Test treatment groups to untreated with significant genes highlighted in red. B & C) PCA variable correlation plots.

**Supplemental Figure 5:** LDH cytotoxicity, pneumolysin activity and *IL-11 RT-PCR with oral commensals.* A) % Cytotoxicity (LDH release) from A549 supernatants 2 hrs post-challenge with TIGR4 (MOI 20), 6B (MOI 20) or uninfected ± indicated treatments (n=4; 2-3 technicals per replicate). UT= untreated. Tukey box and whisker plot, all data was analyzed by One way-ANOVA with Tukey's multiple comparison post-hoc test, **= pV≤0.01, ***= pV≤0.001, ns=not significant. B) Hemolytic activity of A549 cell culture supernatants ± 10 µM GSK-J4 2 hrs post-infection with either 6B (MOI 20) or TIGR4 (MOI 20) with uninfected control. Bar graph ± Std. analyzed by One way-ANOVA with Tukey's multiple comparison post-hoc test, ns=not significant. C) A549 cell protein lysates 2 hrs post-infection, at an MOI of 20, probed for Pneumolysin. Relative band intensity quantified with representative blot images. ΔPlyTIGR is a control. Scatter plot of all replicates with mean denoted in red. Data analyzed by Student's T-Test, ns=not significant. D) Total RNA harvested from immortalized gingival keratinocytes 21 hrs post-challenge with *S. gordonii*, *S. sanguinis*, *S. oralis*, *E. corrodens* or *F. nucleatum* at an MOI of 100. Transcript levels for *IL-11* determined by RT-PCR and represented as relative expression to uninfected (n=1).

**Supplemental Figure 6:** *TIGR4 and 6B bacterial inoculates from animal challenges.* A) 6B and TIGR4 ± GSK-J4 with matched DMSO 24 hrs post-infection bacterial burden from the nasal lavage (NL), bronchoalveolar lavage fluid (BALF), lungs, and spleen of infected animals (n=9). DMSO TIGR4 white, GSK-J4 TIGR4 gray, DMSO 6B light blue, GSK-J4 6B dark blue. Tukey box and whisker plot with dots representing outliers. One way-ANOVA non-parametric Kruskal-Wallis with Dunn's multiple comparison post-hoc test, *= pV≤0.05, **= pV≤0.01, ***= pV≤0.001, ns=not significant. CFU=colony forming unit. Dotted lines =Limit of detection (LD). LD for each organ: NL (50 CFU); BALF, Lung and Spleen (1000 CFU). B) TIGR4 and 6B inoculums for animal intranasal challenge model in DMSO (vehicle control) or 5 mM GSK-J4 (n=3). Conventional CFU enumeration shows no significant difference. Analyzed by One way-ANOVA with Tukey's multiple comparison post-hoc test, ns=not significant. C) TIGR4 ± IL-11 inoculums for animal intranasal challenge model (n=3). Conventional CFU enumeration shows no significant difference. Bar graph ± Std. Analyzed by Student's T-Test, ns=not significant. D) Absorbance ($OD_{600}$) growth curve for mid-log TIGR4 ± IL-11 inoculums for 2 hrs (n=3). No significant difference in growth observed. Mean (dot) ± Std. Analyzed by Student's T-Test at each 20 min interval, ns=not significant.

**Sup. Figure 1**

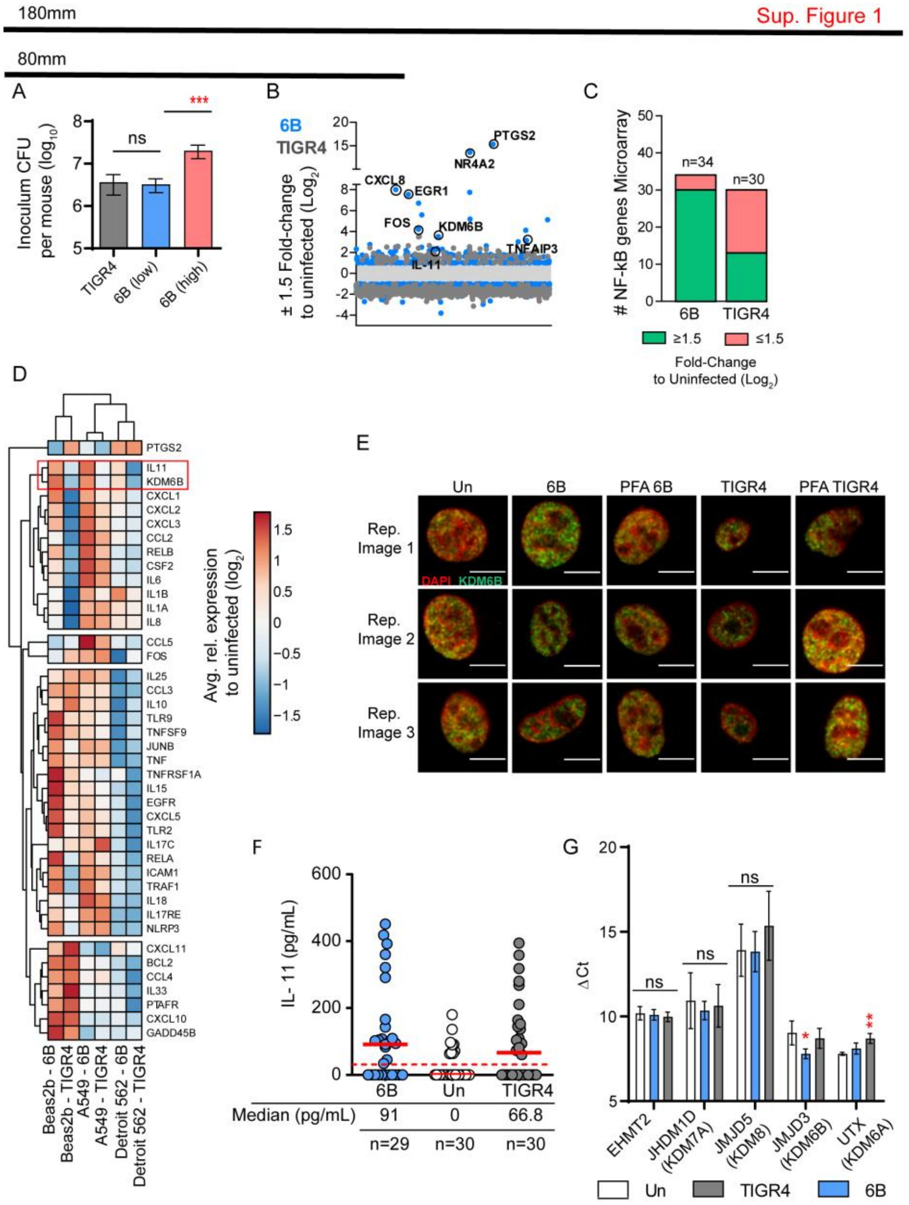


1 Metsalu, T. & Vilo, J. ClustVis: a web tool for visualizing clustering of multivariate data using Principal Component Analysis and heatmap. *Nucleic acids research* **43**, W566-W570, doi:10.1093/nar/gkv468 (2015).

**Sup. Figure 2**

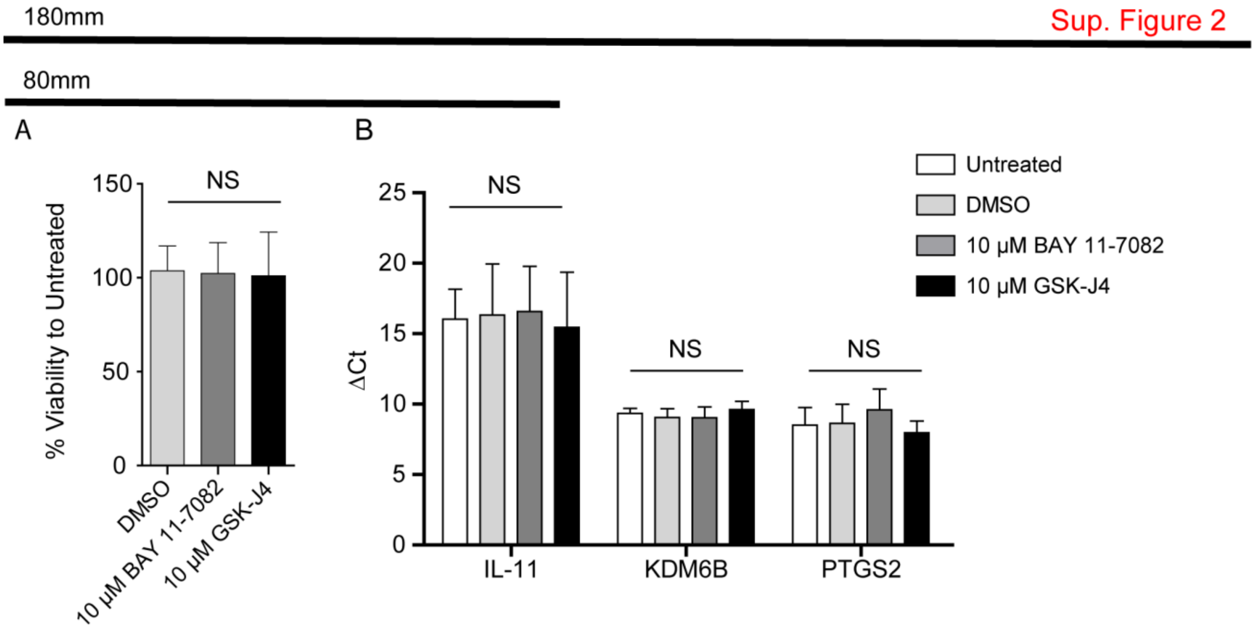
180mm
Sup. Figure 2
80mm
A
B
NS
150
100
50
0
% Viability to Untreated
DMSO
10 μM BAY 11-7082
10 μM GSK-J4
25
20
15
10
5
0
ΔCt
IL-11
KDM6B
PTGS2
Untreated
DMSO
10 μM BAY 11-7082
10 μM GSK-J4

**Sup. Figure 3**

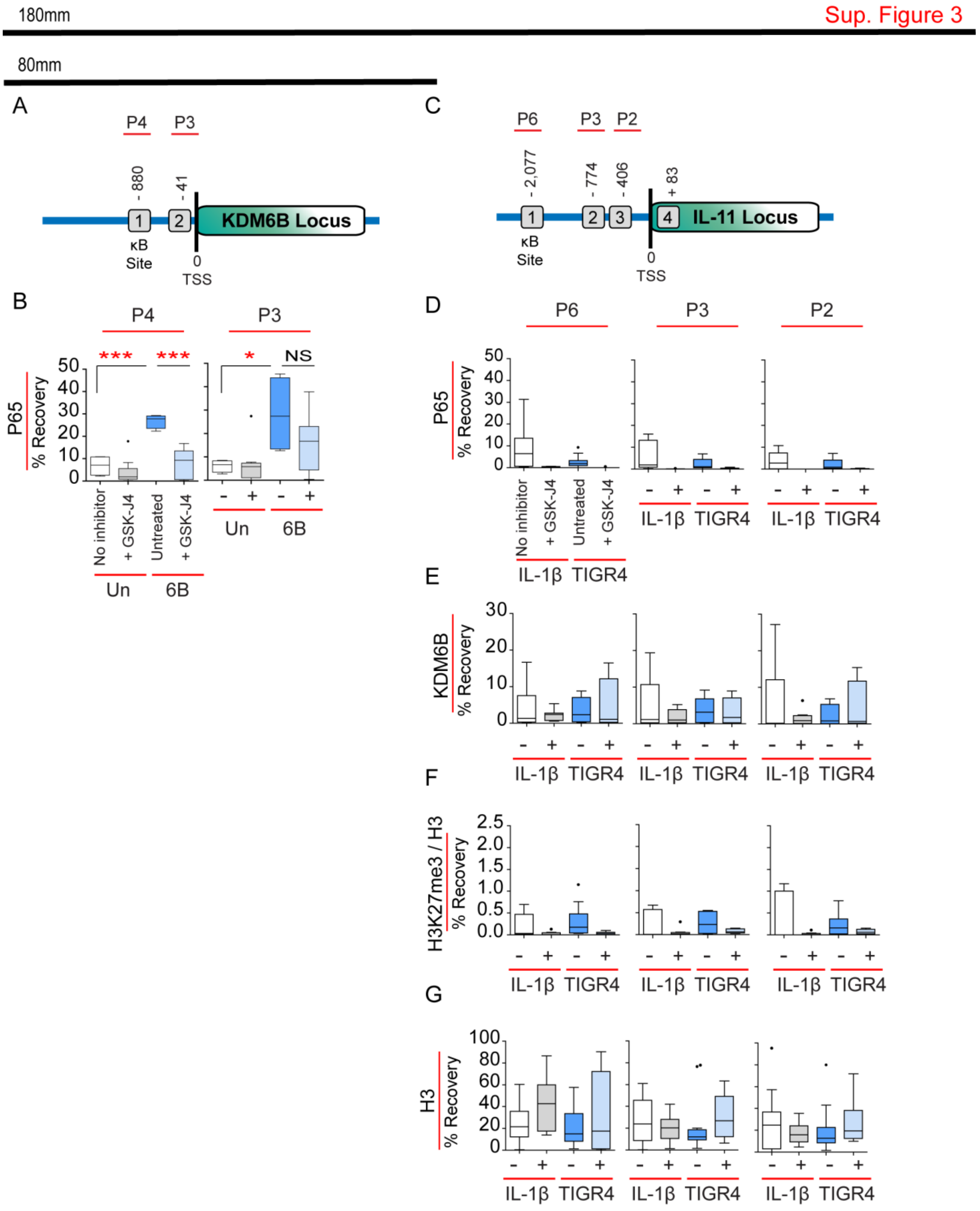

180mm
Sup. Figure 3
80mm
A
P4
P3
- 880
- 41
1
2
KDM6B Locus
κB
Site
0
TSS
C
P6
P3
P2
- 2,077
- 774
- 406
+ 83
1
2
3
4
IL-11 Locus
κB
Site
0
TSS
B
P4
P3
P65
% Recovery
***
***
*
NS
No inhibitor
+ GSK-J4
Untreated
+ GSK-J4
Un
6B
D
P6
P3
P2
IL-1β
TIGR4
E
KDM6B
% Recovery
F
H3K27me3 / H3
% Recovery
G
H3
% Recovery

**Sup. Figure 4**

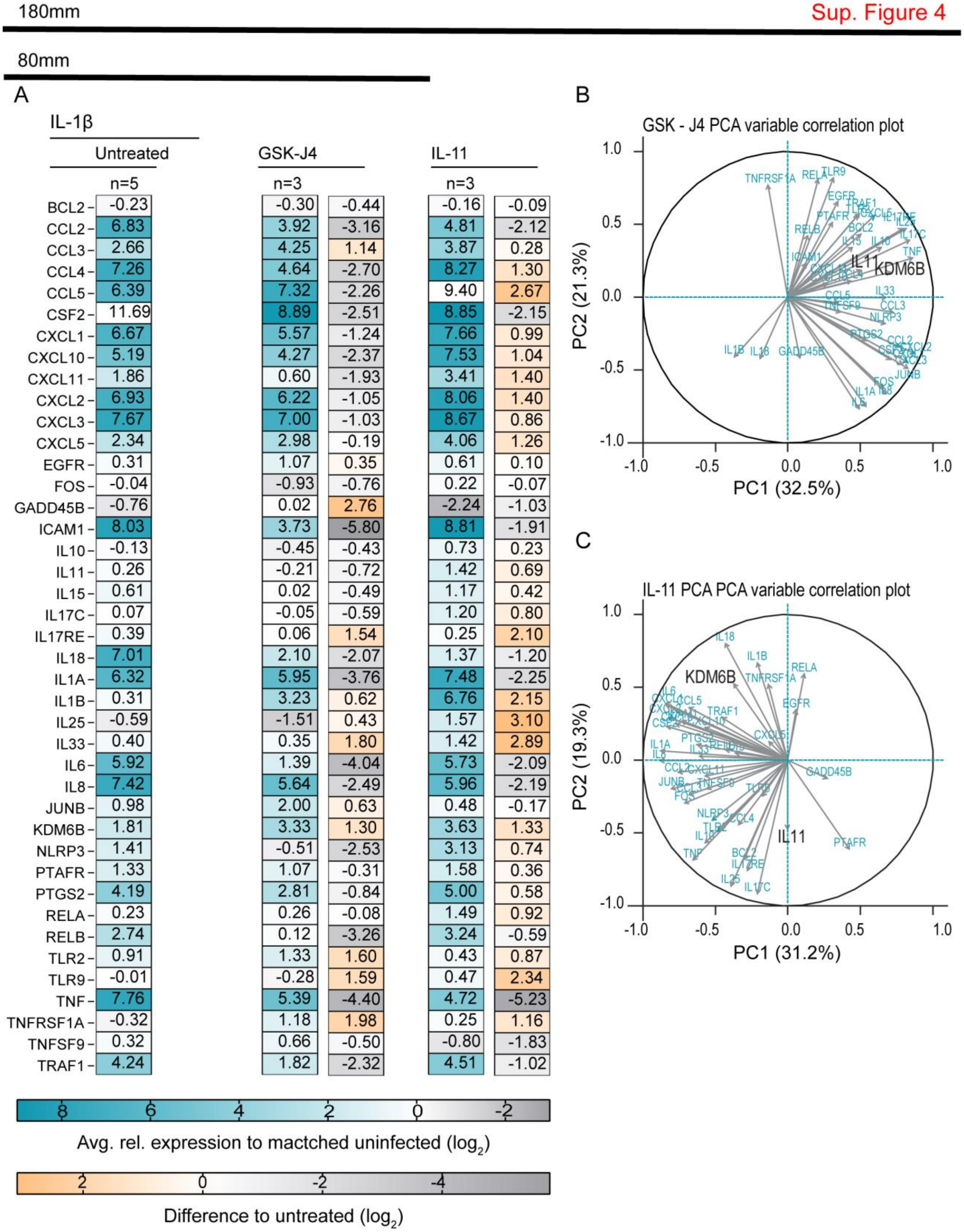


| Gene | IL-1β Untreated (n=5) | GSK-J4 (n=3) | GSK-J4 difference | IL-11 (n=3) | IL-11 difference |
|---|---|---|---|---|---|
| BCL2 | -0.23 | -0.30 | -0.44 | -0.16 | -0.09 |
| CCL2 | 6.83 | 3.92 | -3.16 | 4.81 | -2.12 |
| CCL3 | 2.66 | 4.25 | 1.14 | 3.87 | 0.28 |
| CCL4 | 7.26 | 4.64 | -2.70 | 8.27 | 1.30 |
| CCL5 | 6.39 | 7.32 | -2.26 | 9.40 | 2.67 |
| CSF2 | 11.69 | 8.89 | -2.51 | 8.85 | -2.15 |
| CXCL1 | 6.67 | 5.57 | -1.24 | 7.66 | 0.99 |
| CXCL10 | 5.19 | 4.27 | -2.37 | 7.53 | 1.04 |
| CXCL11 | 1.86 | 0.60 | -1.93 | 3.41 | 1.40 |
| CXCL2 | 6.93 | 6.22 | -1.05 | 8.06 | 1.40 |
| CXCL3 | 7.67 | 7.00 | -1.03 | 8.67 | 0.86 |
| CXCL5 | 2.34 | 2.98 | -0.19 | 4.06 | 1.26 |
| EGFR | 0.31 | 1.07 | 0.35 | 0.61 | 0.10 |
| FOS | -0.04 | -0.93 | -0.76 | 0.22 | -0.07 |
| GADD45B | -0.76 | 0.02 | 2.76 | -2.24 | -1.03 |
| ICAM1 | 8.03 | 3.73 | -5.80 | 8.81 | -1.91 |
| IL10 | -0.13 | -0.45 | -0.43 | 0.73 | 0.23 |
| IL11 | 0.26 | -0.21 | -0.72 | 1.42 | 0.69 |
| IL15 | 0.61 | 0.02 | -0.49 | 1.17 | 0.42 |
| IL17C | 0.07 | -0.05 | -0.59 | 1.20 | 0.80 |
| IL17RE | 0.39 | 0.06 | 1.54 | 0.25 | 2.10 |
| IL18 | 7.01 | 2.10 | -2.07 | 1.37 | -1.20 |
| IL1A | 6.32 | 5.95 | -3.76 | 7.48 | -2.25 |
| IL1B | 0.31 | 3.23 | 0.62 | 6.76 | 2.15 |
| IL25 | -0.59 | -1.51 | 0.43 | 1.57 | 3.10 |
| IL33 | 0.40 | 0.35 | 1.80 | 1.42 | 2.89 |
| IL6 | 5.92 | 1.39 | -4.04 | 5.73 | -2.09 |
| IL8 | 7.42 | 5.64 | -2.49 | 5.96 | -2.19 |
| JUNB | 0.98 | 2.00 | 0.63 | 0.48 | -0.17 |
| KDM6B | 1.81 | 3.33 | 1.30 | 3.63 | 1.33 |
| NLRP3 | 1.41 | -0.51 | -2.53 | 3.13 | 0.74 |
| PTAFR | 1.33 | 1.07 | -0.31 | 1.58 | 0.36 |
| PTGS2 | 4.19 | 2.81 | -0.84 | 5.00 | 0.58 |
| RELA | 0.23 | 0.26 | -0.08 | 1.49 | 0.92 |
| RELB | 2.74 | 0.12 | -3.26 | 3.24 | -0.59 |
| TLR2 | 0.91 | 1.33 | 1.60 | 0.43 | 0.87 |
| TLR9 | -0.01 | -0.28 | 1.59 | 0.47 | 2.34 |
| TNF | 7.76 | 5.39 | -4.40 | 4.72 | -5.23 |
| TNFRSF1A | -0.32 | 1.18 | 1.98 | 0.25 | 1.16 |
| TNFSF9 | 0.32 | 0.66 | -0.50 | -0.80 | -1.83 |
| TRAF1 | 4.24 | 1.82 | -2.32 | 4.51 | -1.02 |

**Sup. Figure 5**

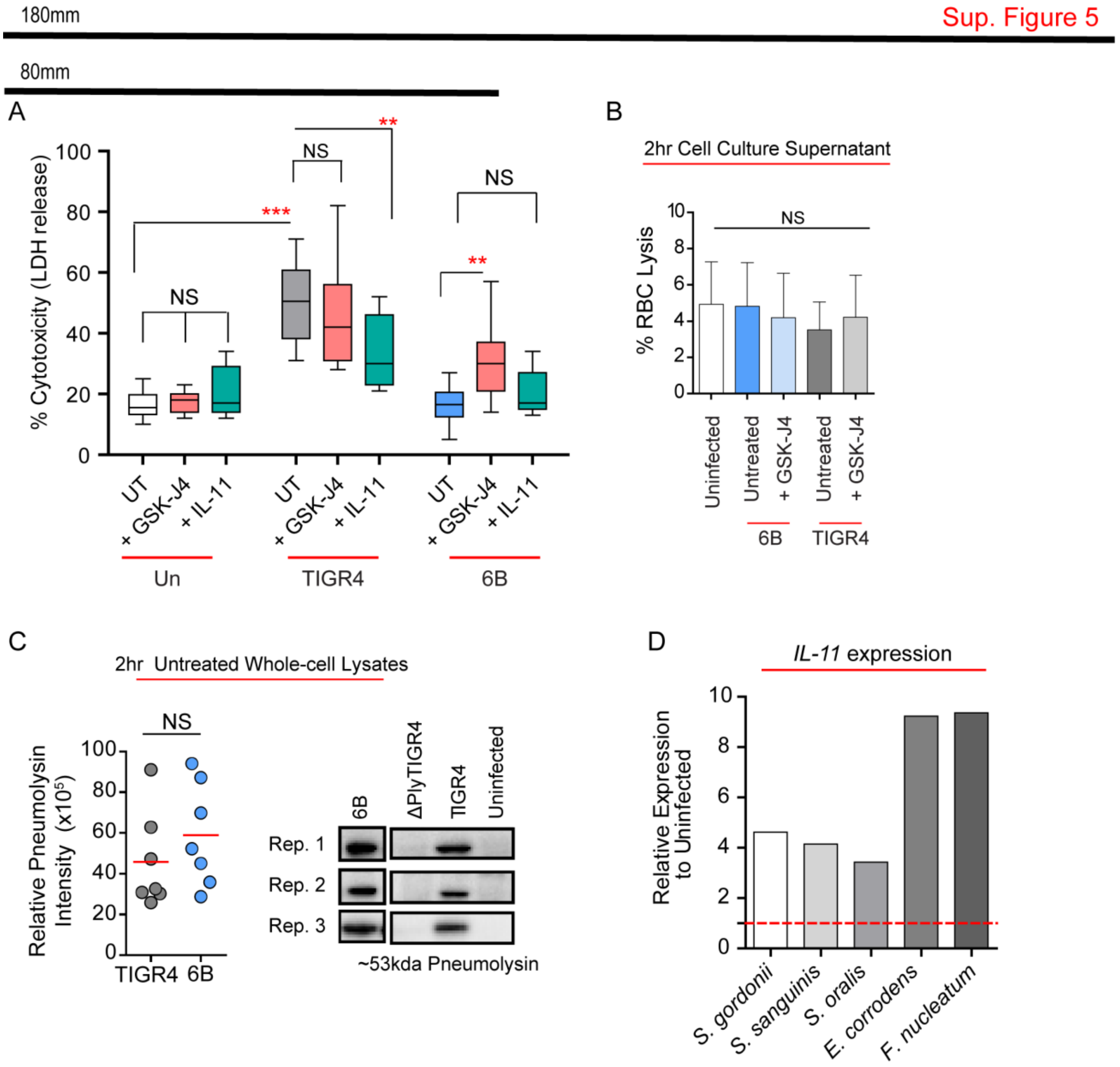
180mm
Sup. Figure 5
80mm
A
% Cytotoxicity (LDH release)
100
80
60
40
20
0
***
**
NS
NS
NS
**
UT
+ GSK-J4
+ IL-11
UT
+ GSK-J4
+ IL-11
UT
+ GSK-J4
+ IL-11
Un
TIGR4
6B
B
2hr Cell Culture Supernatant
% RBC Lysis
10
8
6
4
2
0
NS
Uninfected
Untreated
+ GSK-J4
Untreated
+ GSK-J4
6B
TIGR4
C
2hr Untreated Whole-cell Lysates
NS
Relative Pneumolysin Intensity (x10^5)
100
80
60
40
20
0
TIGR4
6B
6B
ΔPlyTIGR4
TIGR4
Uninfected
Rep. 1
Rep. 2
Rep. 3
~53kda Pneumolysin
D
IL-11 expression
Relative Expression to Uninfected
10
8
6
4
2
0
S. gordonii
S. sanguinis
S. oralis
E. corrodens
F. nucleatum

**Sup. Figure 6**

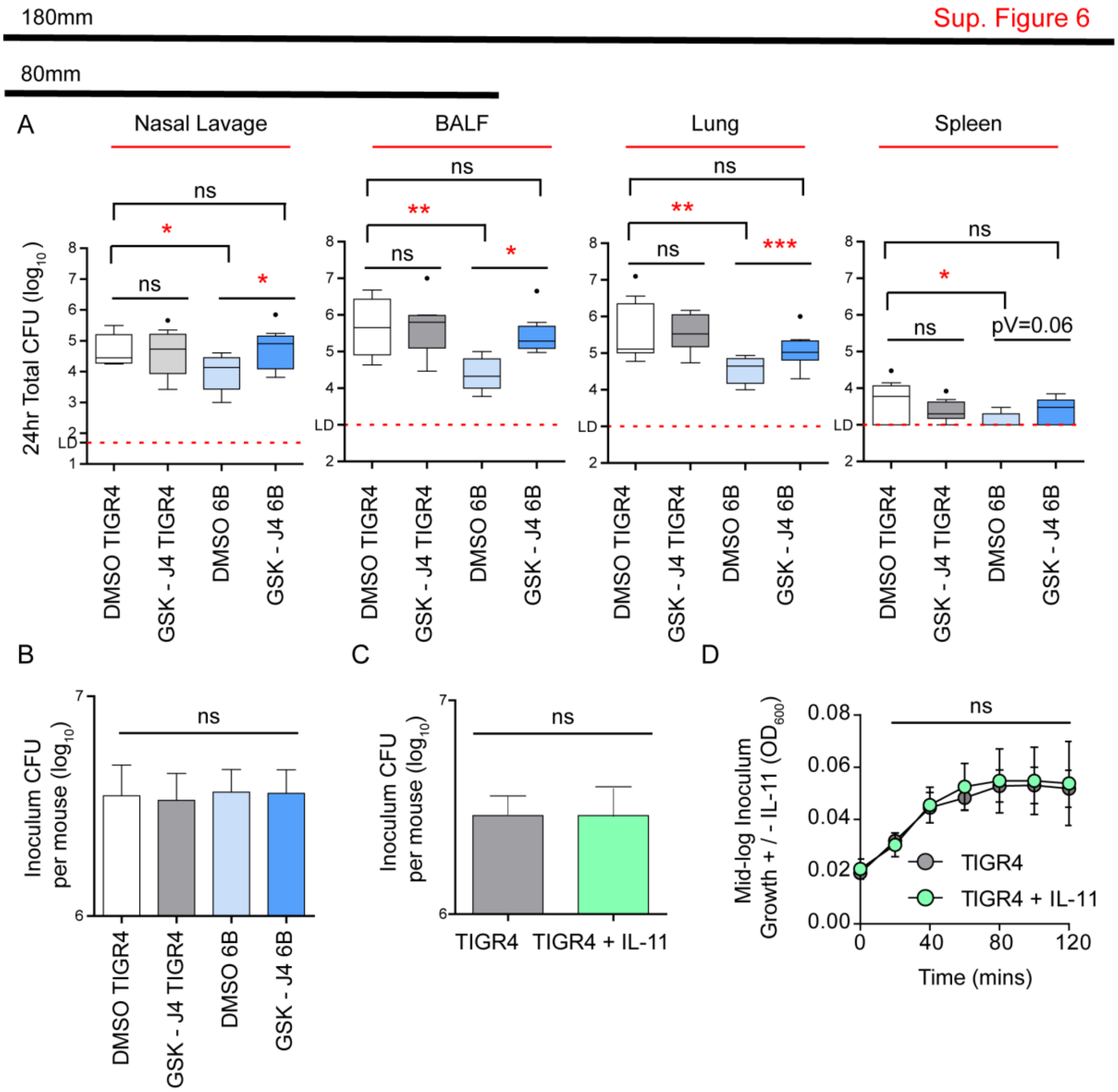
180mm
Sup. Figure 6
80mm
A
Nasal Lavage
BALF
Lung
Spleen
24hr Total CFU (log10)
ns
*
**
***
pV=0.06
LD
DMSO TIGR4
GSK - J4 TIGR4
DMSO 6B
GSK - J4 6B
B
Inoculum CFU per mouse (log10)
C
TIGR4
TIGR4 + IL-11
D
Mid-log Inoculum Growth + / - IL-11 (OD600)
Time (mins)
